# An *Engrailed1* enhancer underlies human thermoregulatory evolution

**DOI:** 10.1101/2020.08.22.262659

**Authors:** Daniel Aldea, Yuji Atsuta, Blerina Kokalari, Stephen Schaffner, Bailey Warder, Yana Kamberov

## Abstract

Humans rely on sweating to cool off and have the highest eccrine sweat gland density among mammals. We investigated whether altered regulation of the *Engrailed 1* (*EN1*) gene, the levels of which are critical for patterning eccrine glands during development, could underlie the evolution of this defining human trait. First, we identify five *EN1* candidate enhancers (ECEs) using comparative genomics and validation of enhancer activity in mouse skin. The human ortholog of one ECE, hECE18, contains multiple derived substitutions that together dramatically increase the activity of this enhancer in keratinocytes. Targeted repression of hECE18 reduces *EN1* expression in human keratinocytes, indicating hECE18 upregulates *EN1* in this context. Finally, we find that hECE18 increases ectodermal *En1* in a humanized knock-in mouse to increase eccrine gland number. Our study uncovers a genetic basis for the evolution of one of the most singular human adaptations and implicates the recurrent mutation of a single enhancer as a novel mechanism for evolutionary change.

## INTRODUCTION

Because of their furlessness, humans are often called the “naked ape”, but humans are also the “sweaty ape” (Newman, 1970). Sweating is the major mechanism by which humans dissipate body heat: evaporative cooling that occurs when water (sweat) is vaporized on the skin (Kuno, 1956). Humans can secrete upwards of 1L of sweat per hour, and thermally induced sweat rates in humans are reported to be four to ten times higher than those of chimpanzees (Folk and Semken, 1991; Hiley, 1976). Because fur reduces airflow over the skin (Al-Ramamneh et al., 2011), human furlessness is thought to be an adaptation that enhances sweat evaporation (Carrier, 1984; Lieberman, 2015; Montagna, 1963, 1972). A second critical adaptation that also underlies humans’ exceptional sweating capabilities is a high density of eccrine sweat glands in the skin (Folk and Semken, 1991; Kamberov et al., 2018; Montagna, 1963, 1972). Eccrine glands secrete the water humans vaporize for evaporative cooling (Kuno, 1956) and their importance is underscored by the risk of hyperthermia in individuals with reduced numbers of these organs (Wright et al., 1993). Eccrine glands are the predominant appendages of human skin, with densities exceeding 200 glands/cm^2^ in regions such as the face (Kamberov et al., 2018). In comparison, eccrine gland densities of macaques and chimpanzees are on average ten times lower than that of humans (Kamberov et al., 2018). How and when humans diverged from other primates to evolve this dramatically elaborated eccrine gland density is unknown.

The evolution of humans’ high eccrine gland density required modification to the program that controls the number of eccrine glands specified during development. Much of our understanding of this developmental program comes from studies of eccrine gland formation in mice. Mice retain the ancestral condition of having eccrine glands only in the volar (palmar/plantar) skin of the paws, where eccrine gland secretions regulate frictional contact with underlying surfaces (Adelman et al., 1975). We and others have previously shown that the transcription factor Engrailed 1 (*En1*) promotes the specification of eccrine glands in elevations of the volar skin called footpads, and also in the intervening interfootpad space (IFP) of the mouse paw (Aldea et al., 2019; Kamberov et al., 2015; Loomis et al., 1996; Lu et al., 2016). In this context, *En1* is expressed throughout the basal keratinocyte layer of developing mouse volar skin and is specifically upregulated in eccrine gland placodes of the IFP (Kamberov et al., 2015). *En1* knock-out mice fail to form eccrine glands and reducing *En1* levels in this species results in a dose-dependent decrease in volar eccrine gland number (Aldea et al., 2019; Kamberov et al., 2015; Loomis et al., 1996). In fact, intrinsic differences in the *cis*-regulation of *En1* are a primary cause of the four-fold greater abundance of IFP eccrine glands in FVB/N as compared to C57BL/6N mouse strains (Kamberov et al., 2015). Strikingly, *EN1* is upregulated in human fetal ectoderm co-incident with the onset of eccrine gland specification (Lu et al., 2016). In light of these findings, we investigated whether evolutionary changes leading to increased ectodermal *EN1* during development could be an underlying mechanism for the adaptive increase in human eccrine gland density.

## RESULTS

### Identification of candidate *Engrailed 1* enhancers in the skin

The *EN1* coding sequence is highly conserved among primates and most evolutionarily-relevant changes are thought to reside in regulatory DNA (King and Wilson, 1975). Therefore, to determine if humans have altered *EN1* regulation, we first screened for regulatory elements, or enhancers, with the potential to control ectodermal *EN1* during eccrine gland development. Direct interrogation of human developmental enhancers in this context is confounded by the fact that there are no *in vitro* systems that recapitulate human eccrine gland development, nor indeed of any species, and eccrine gland placodes begin to form during the second trimester in humans (William Montagna and Paul F. Parakhal, 1974). Accordingly, we used a candidate-based screen to identify putative enhancers. In general, patterns of ectodermal *EN1* expression are similar in body regions where eccrine glands form across mammals (Kamberov et al., 2015; Lu et al., 2016; Mainguy et al., 1999), therefore we sought to identify putative EN1 enhancers based on evidence of evolutionary sequence conservation (Figure 1A, Methods). We restricted our screen to non-coding DNA withing the genomic interval spanned by the *EN1* topologically associated domain (TAD) as defined by Hi-C in normal human cultured keratinocytes (Chr2: 118000000-118880000 (hg38)) since studies on the compartmentalization of the mammalian genome suggest that elements controlling expression of *EN1*, or indeed any locus, are likely to be located within this smaller interval (Dixon et al., 2012, 2015). Importantly, the genomic interval spanned by the human *EN1* TAD is in a region of conserved synteny to the mapped mouse *En1* TAD (Chr1:120583423-121463423 (mm10)) (Dixon et al., 2012). Using phastCons (Siepel et al., 2005), we called 209 evolutionarily conserved elements within the *EN1* TAD that were at least 50 base pairs (bps) in length and ≥98% conserved across the phastCons60way placental mammals dataset (Figure S1A and Methods). We generated a list of 41 priority conserved elements based on overlap with published datasets of genomic regions that contain epigenomic marks suggestive of enhancer presence from both mouse and human datasets and also with published catalogs of human genomic regions that show evidence of accelerated evolution (Capra et al., 2013; Pollard et al., 2006; Prabhakar et al., 2006). Using each priority element as a kernel, we expanded these genomic regions to 1000bp-1500bp because known enhancers tend to be hundreds of base pairs long, collapsing our list to 23 *Engrailed 1* candidate enhancers (ECEs) for functional testing (Figure 1B and S1B).

**Figure 1.**
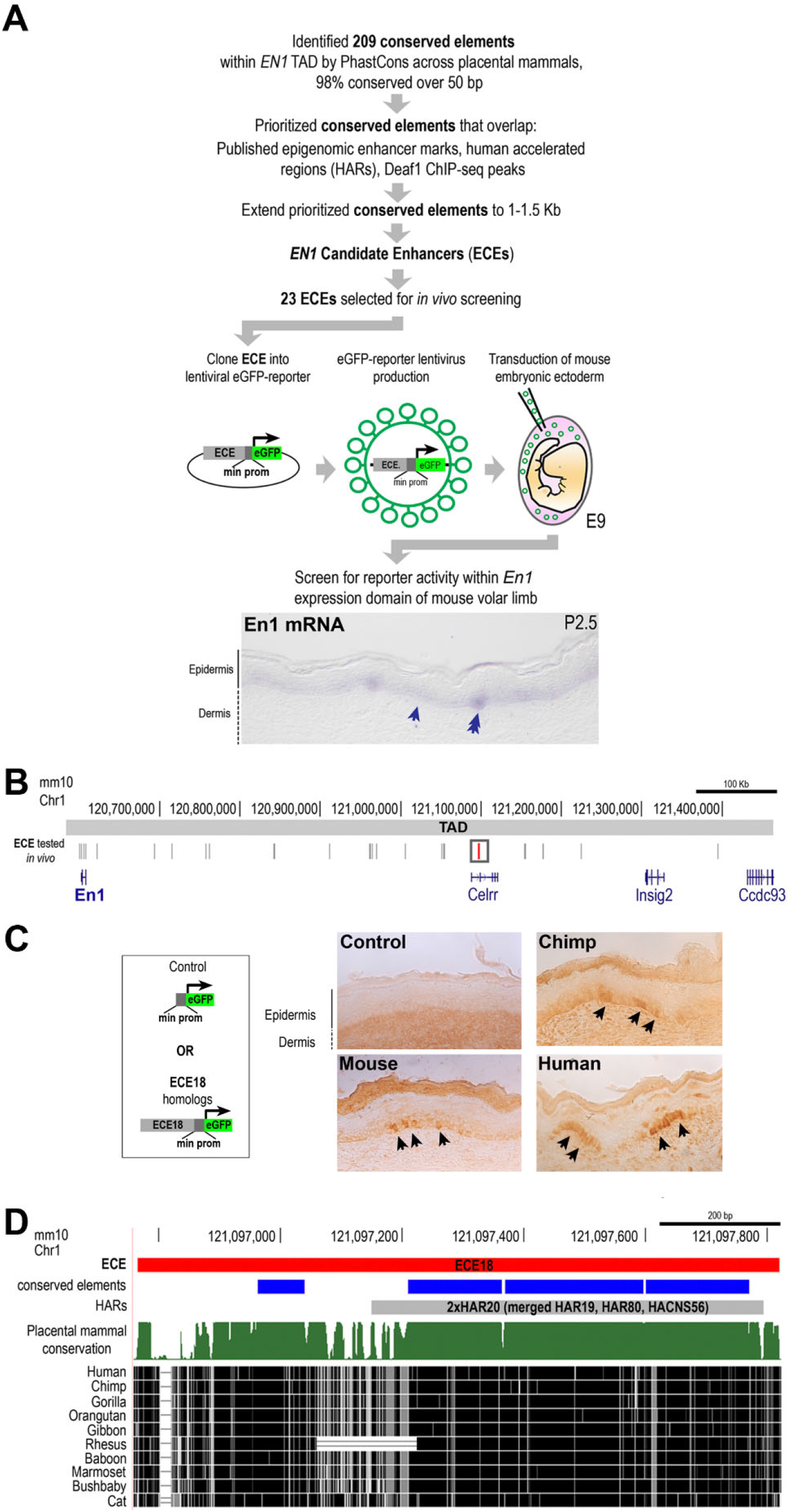
Identification of *Engrailed 1* candidate enhancer ECE18. **(A)** Strategy to identify putative developmental enhancers of *En1*. *in situ* hybridization for *En1* mRNA (purple) in mouse volar limb skin at P2.5. Basal keratinocyte layer (arrow) and eccrine placode (double arrow). **(B)** Location in mm10 of *En1* candidate enhancers (ECEs) tested *in vivo* (vertical grey lines). ECE18 is in red and boxed. **(C)** Representative images of mouse, chimpanzee and human ECE18 transgenic P2.5 volar limb stained with anti-GFP antibody. eGFP (black arrow) is visualized using HRP-DAB coupled immunohistochemistry. **(D)** Sequence alignment and evolutionarily features of ECE18, which contains four conserved elements by phastCons (blue lines) and overlaps the human accelerated region 2xHAR20.

We evaluated the potential of ECEs to act as ectodermal *En1* enhancers by analyzing whether each element was able to activate reporter expression in the *En1*-postive domain of developing mouse volar skin. To this end, we used lentiviruses encoding the mouse ortholog of each ECE upstream of a minimal promoter and eGFP reporter cassette to stably transduce embryonic mouse ectoderm and generate skin-specific transgenic ECE reporter mice (Figure 1A and Methods) (Beronja et al., 2010; Emerson and Cepko, 2011; Wang et al., 2014).

Of the 23 tested ECEs, five consistently produced eGFP-positive clones in the basal (*En1*-positive) keratinocyte layer of mouse volar skin on post-natal day (P) 2.5 (Figure S1B). At this stage, *En1* is critical for specifying the number of eccrine glands in the IFP and is expressed throughout basal keratinocytes of the distal volar skin and focally upregulated in IFP eccrine gland placodes (Figure 1A) (Kamberov et al., 2015). The five positive ECEs consisted of a fragment of the *En1* promoter (ECE2, Chr1: 120601976-120602486 (mm10)), and four elements located downstream of *En1* (ECE8, Chr1: 120756823-120757766; ECE18, Chr1: 121096764-121097826; ECE20, Chr1: 121176848-121178300; ECE23, Chr1: 121394405-121395702 (mm10)) (Figure 1C and S1B). All positive ECEs also produced positive clones within the basal ectoderm of the volar footpads and differentiating eccrine glands therein, indicating these ECEs can function at later stages of eccrine gland development (Figure S2A). We noted that ECE2, ECE18 and ECE20 transgenic mice also had eGFP-positive clones in the dorsal limb skin, which is outside of the *En1-*positive domain in mice (Figure S2B). This is intriguing given that *EN1* expression in the basal ectoderm is expanded to the non-volar skin in humans.

### Enhanced activity of the human ECE18 ortholog

Since our goal was to identify ECEs with the potential to explain a human-specific phenotype, we focused our subsequent analyses on ECE18. This element contains the highest number of derived human mutations. Moreover, ECE18 overlaps the HACNS56, HAR19 and HAR80 human accelerated regions (collectively named 2xHAR20) which are class of genomic elements that are highly conserved across vertebrates but are exceptionally diverged in humans, suggestive of evolutionary importance in our species (Capra et al., 2013; Pollard et al., 2006; Prabhakar et al., 2006) (Figure 1D and Figure S3A).

Consistent with having ectodermal enhancer capabilities during development, the human and chimpanzee orthologs of ECE18 (hECE18 and cECE18, respectively) induced eGFP-positive clones in limb skin in a pattern identical to that of mouse ECE18 (Figure 1C and S2A, B). However, while all tested ECE18 orthologs behaved similarly in the mouse transgenic assay, comparison of the activity of ECE18 orthologs in quantitative luciferase reporter assays in cultured human keratinocytes revealed dramatic differences in the potency of this enhancer between species (Figure 2A and S3B, C and Methods). When cloned upstream of a minimal promoter driving luciferase expression, we found that anthropoid ECE18 orthologs produced the greatest fold luciferase induction, while orthologs from species outside this primate infraorder induced little to no luciferase activity (Figure 2A). Among tested orthologs, hECE18 was the most potent enhancer, producing on average a 13-fold increase in luciferase over control. Within catarrhines, the group consisting of Old World Monkeys (OWM’s) and apes, hECE18 was 2.6-fold and 1.4-fold higher relative to chimpanzee and macaque orthologs, respectively (Figure 2A).

**Figure 2.**
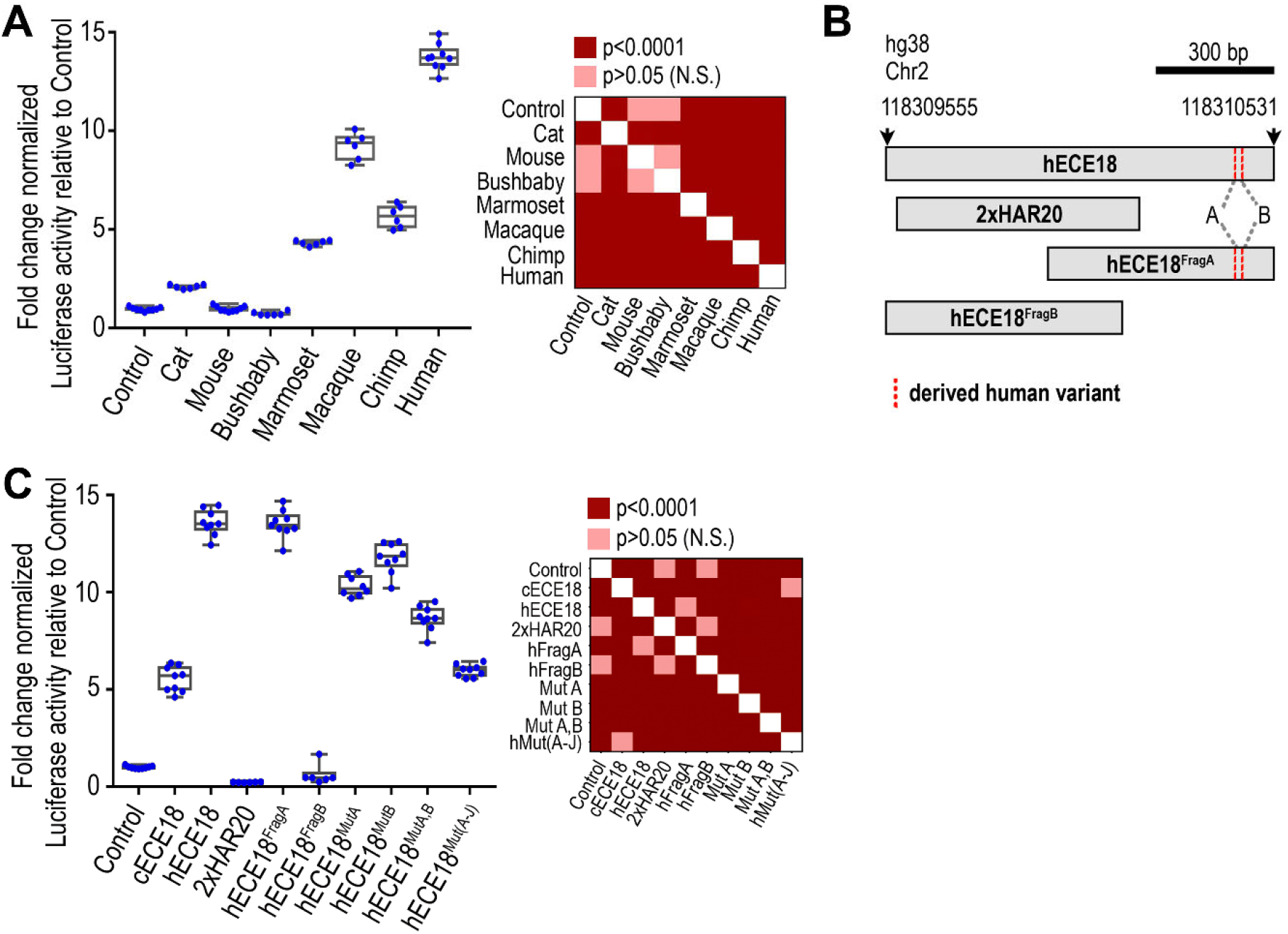
Recurrent evolution of human ECE18 produced human-specific gains in enhancer activity. **(A)** Activity of ECE18 orthologs in human keratinocytes. Fold change normalized luciferase activity is plotted. **(B,C)** Regional dissection and mapping of sequence variants underlying human ECE18 (hECE18) activity. Derived human bases at A and B are in red (Chr2:118,310,426 and Chr2:118,310,438, respectively hg38). Chimpanzee ECE18 (cECE18). Mutagenized hECE18 (Mut) with substitution of indicated derived human base(s) to ancestral ape base(s). All ten derived human substitutions are mutated to ancestral ape in hECE18^MutA-J^. Normalized firefly luciferase activity is plotted as the fold change relative to Control (empty reporter vector). Firefly luciferase values normalized to Renilla luminescence. Dots represent an individual biological replicate. Median (line), box (bounds 25%-75%) and whiskers (min and max) plotted. Significance by one-way ANOVA. Tukey-adjusted *P*-values are reported. Assays performed in human GMA24F1A keratinocytes.

### Recurrent mutation of ECE18 during human evolution

That ECE18 activity is higher in humans, suggested that human-specific sequences may underlie the potency of this enhancer in our species. Accordingly, we tested the 2xHAR20 genomic fragment that contains the highest number of derived human substitutions in hECE18 for enhancer activity (Figure 1D and S3A). 2xHAR20 did not induce reporter expression in cultured keratinocytes or in mouse transgenic assays, indicating that this region is not sufficient for enhancer activity (Figure 2B, C). Thus, we split hECE18 into two partially overlapping fragments: hECE18^FragB^ (hg38 Chr2: 118309555-118310154bp) and hECE18^FragA^ (hg38 Chr2: 118309932-118310531bp) (Figure 2B and S3C). hECE18^FragB^ did not induce reporter expression *in vivo* or *in vitro* (Figure 2C). In contrast, hECE18^FragA^ recapitulated both the quantitative activity and spatial/temporal activity of full-length hECE18 in cultured human keratinocytes and P2.5 mouse transgenic skin, respectively (Figure 2C and S3C-E). Of note hECE18^FragA^ is conserved to Neanderthals and Denisovans indicating equivalent levels of enhancer activity in these hominins. ECE18^FragA^ orthologs from non-human anthropoids also recapitulated the quantitative activity of the respective full-length elements (Figure 3E). As with full length hECE18, hECE18^FragA^ produced the same fold-change increase in luciferase relative to tested non-human primate orthologs consistent with the idea that this region is sufficient to explain enhancer activity in all species (Figure 2C and S3E).

**Figure 3.**
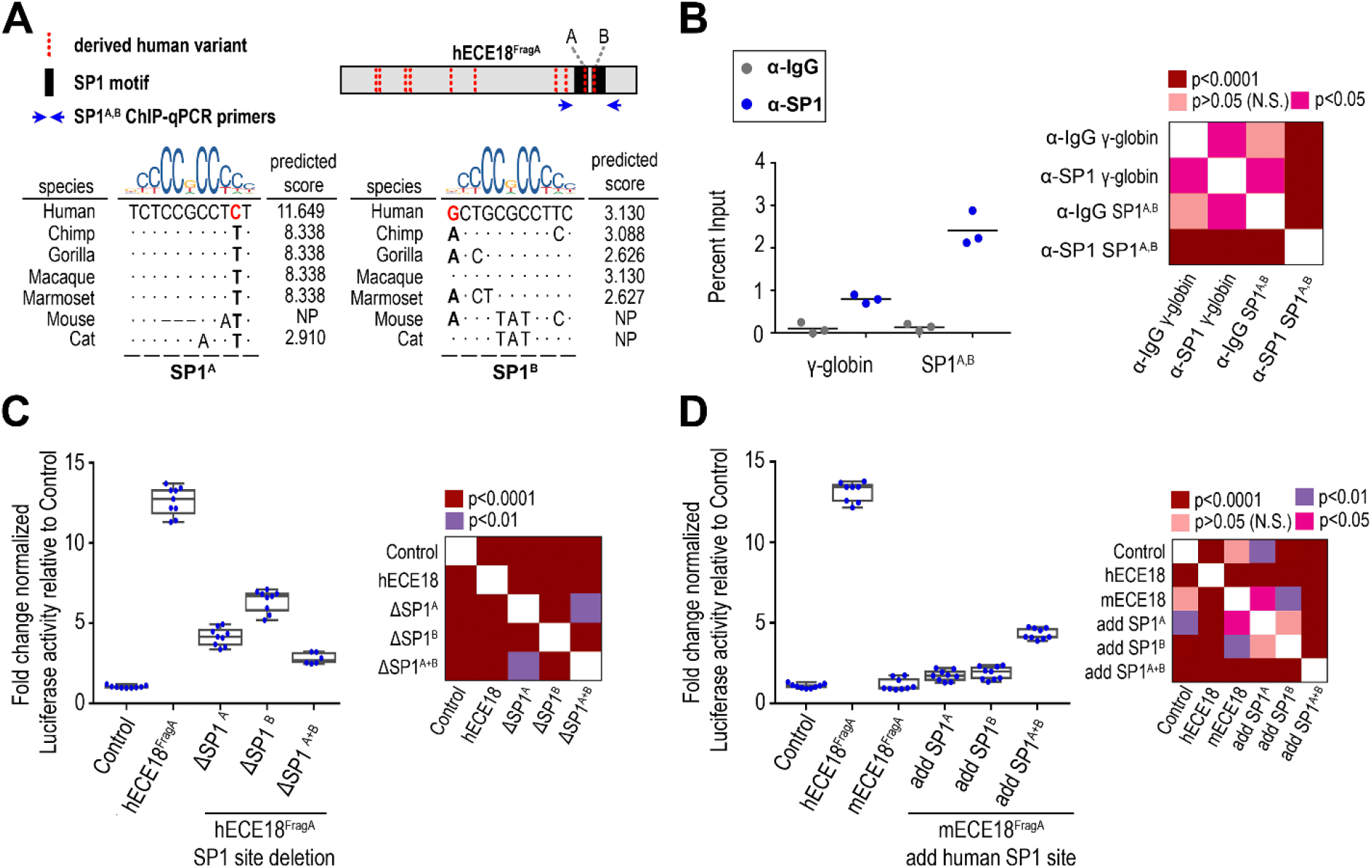
Variation at SP1 binding motifs underlies evolutionary changes in ECE18 activity. **(A)** Alignment and predicted SP1 binding affinities for SP1 at SP1^A^ and SP1^B^. Derived human bases at A and B are in red. Dots indicate identity to human base. Blue arrows (SP1^A,B^ ChIP-qPCR primers). **(B)** Enrichment of SP1 by ChIP-qPCR at hECE18 interval containing SP1^A^ and SP1^B^ motifs (SP1^A, B^) in human keratinocytes. Human γ-globin (HBG2) promoter is used as a negative control and IgG or SP1 enrichment over the input for each set of primers is shown. Mean enrichment across three independent experiments (line). **(C)** Fold change luciferase induction by hECE18^FragA^ upon deletion of SP1^A^ and SP1^B^. **(D)** Fold change luciferase induction by mouse ECE18^FragA^ (mECE18 ECE18^FragA^) in human keratinocytes upon knock-in of human SP1^A^ and SP1^B^ motifs. Normalized firefly luciferase activity is plotted as the fold change relative to Control (empty reporter vector). Firefly luciferase values normalized to Renilla luminescence. Dots represent an individual biological replicate. Median (line), box (bounds 25%-75%) and whiskers (min and max) plotted. Significance by one-way ANOVA. Tukey-adjusted *P*-values are reported. Assays performed in human GMA24F1A keratinocytes.

We found that replacing even a portion of the genomic interval spanned by hECE18^FragA^ in hECE18 with chimpanzee sequence was sufficient to reduce the activity of the enhancer in luciferase assays, indicating the importance of human sequence changes within this enhancer (Figure S3F). hECE18^FragA^ contains 12 derived human mutations relative to other apes, including two single nucleotide insertions and 10 single nucleotide substitutions (called here sites A-J) (Figure 3G). Deletion of the two derived insertions or individual mutation of substitutions C-J to the ancestral ape base in hECE18 did not significantly affect enhancer activity (Figure S3H, I). However, mutation of the derived human nucleotides at A and B, alone or in combination, to the ancestral ape bases resulted in a consistent reduction of enhancer activity (Figure 2C and S3I). Intriguingly, mutation of all ten derived human substitutions (A-J) in combination to the ancestral bases reduced hECE18 activity to that of non-human apes (Figure 2C and S3I).

These results suggest that the epistatic effects of multiple human-specific variants located within the interval spanned by hECE18^FragA^ are responsible for the increased potency of this enhancer in humans relative to other apes. From an evolutionary perspective, this means that ECE18 was subjected to recurrent mutation over the course of human evolution. It is also worth noting that five of these derived human-specific substitutions (namely C-G) are all located within the region overlapping 2xHAR20. This indicates that while the accelerated region is not sufficient for enhancer activity, its evolution played a role in driving the full magnitude of increase in the potency of the hECE18 enhancer.

### A conserved mechanism for ECE18 evolution

*in silico* motif analysis revealed that sites A and B lie within tandem predicted SP1 binding motifs, which we named SP1^A^ and SP1^B^, respectively (Figure 3A). We confirmed SP1 occupancy by ChIP-qPCR in the region spanned by SP1^A^ and SP1^B^ (Figure 3B). Deleting SP1^A^ or SP1^B^ in hECE18^FragA^ reduced luciferase levels to less than half of the wildtype human enhancer, with loss of both motifs reducing enhancer activity to nearly control levels (Figure 3C).

Overall, anthropoids are predicted to have the highest affinity SP1^A^ and SP1^B^ motifs among mammals, and ECE18 anthropoid orthologs had the highest activity *in vitro* (Figures 2A and 3A). The correlation between higher affinity motifs at SP1^A^ and SP1^B^ and increased ECE18 potency is also observed within anthropoids, with humans having the highest affinity sites among apes and the quantitatively strongest enhancer in keratinocytes (Figure 2A). In line with this, the presence of a human-like SP1^B^ motif in OWMs may help to explain why tested OWM orthologs were more active in luciferase assays than those of non-human apes (Figure 2A and S3E, K).Consistent with a model in which sequence variation at SP1^A/B^ underlies phylogenetic variation in ECE18 potency, the introduction of human SP1^A^ and SP1^B^ motifs into mouse ECE18^FragA^, which is not predicted to bind SP1 at either position, additively increased luciferase induction by the mouse enhancer (Figure 3D).

### ECE18 modulates endogenous *EN1* expression in human keratinocytes

Having identified an enhancer in which human-specific sequence evolution increased activity in keratinocytes, we investigated whether hECE18 regulates *EN1* in this context. We repressed the endogenous hECE18 enhancer in cultured human keratinocytes using a dCas9-KRAB domain fusion and gRNAs targeting the ECE18 genomic interval (Figure 4A). We found that targeting the repressor complex to hECE18 reduced *EN1* mRNA by 40% on average relative to control, demonstrating that hECE18 positively regulates *EN1* expression (Fig.4B).

**Figure 4.**
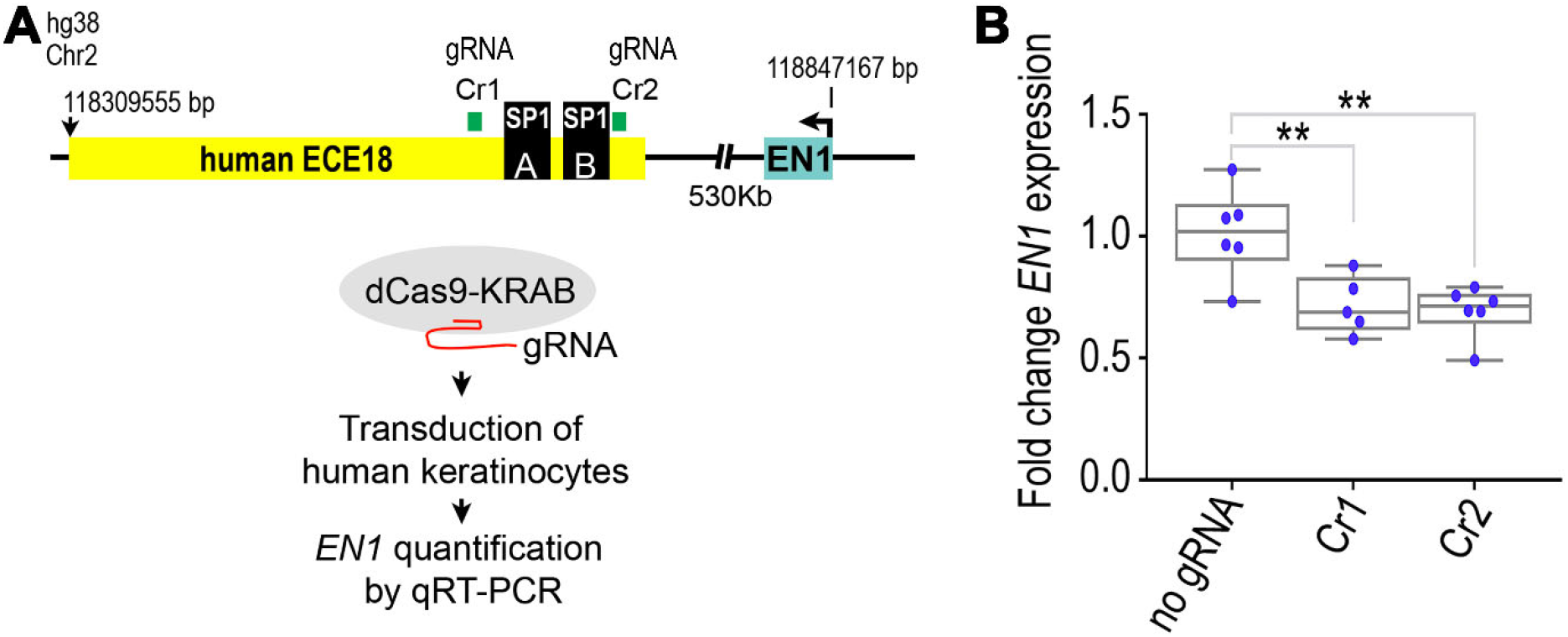
hECE18 positively regulates *EN1* expression in human keratinocytes. **(A)** Overview of strategy to repress hECE18 in human keratinocytes using dCas9-KRAB domain fusion complex. Cr1 and Cr2 guide RNAs target hECE18. Relative genomic positions of *EN1*, SP1^A^ and SP1^B^ motifs are shown. **(B)** Fold change normalized *EN1* expression by qRT-PCR in human GMF24F1A keratinocytes upon dCas9-KRAB repression of hECE18. Fold change calculated relative to dCAS9-KRAB transduction alone. Dots represent an individual biological replicate. Median (line), box (bounds 25%-75%) and whiskers (min and max) plotted. Significance by one-way ANOVA. Tukey-adjusted *P*-values are reported: ** *P*<0.01.

### Human ECE18 upregulates *En1* to promote eccrine gland specification

To determine if hECE18 can function as a developmental enhancer of ectodermal *Engrailed 1*, we turned to the mouse. We first determined if the higher potency of hECE18 could be recapitulated in this model. Consistently, hECE18 produced the greatest fold luciferase induction in cultured mouse primary keratinocytes (Figure 5A). We noted, however, the relative increase over control was 4-fold and 1.5-fold on average by hECE18 and cECE18, respectively, which was markedly lower than observed in human skin cells for both orthologs. This suggests that the relative activity of ECE18 may be attenuated in this system. As in human keratinocytes, mouse ECE18 did not induce luciferase expression significantly. Moreover, in contrast to our findings in repressing hECE18, deletion of the endogenous ECE18 sequence from the mouse genome did not alter ectodermal *En1* expression or eccrine gland number in mouse volar skin (Figure S4). These data show that though ECE18 is not essential for *En1* regulation in mouse skin cells, the human ECE18 ortholog functions in this context.

**Figure 5.**
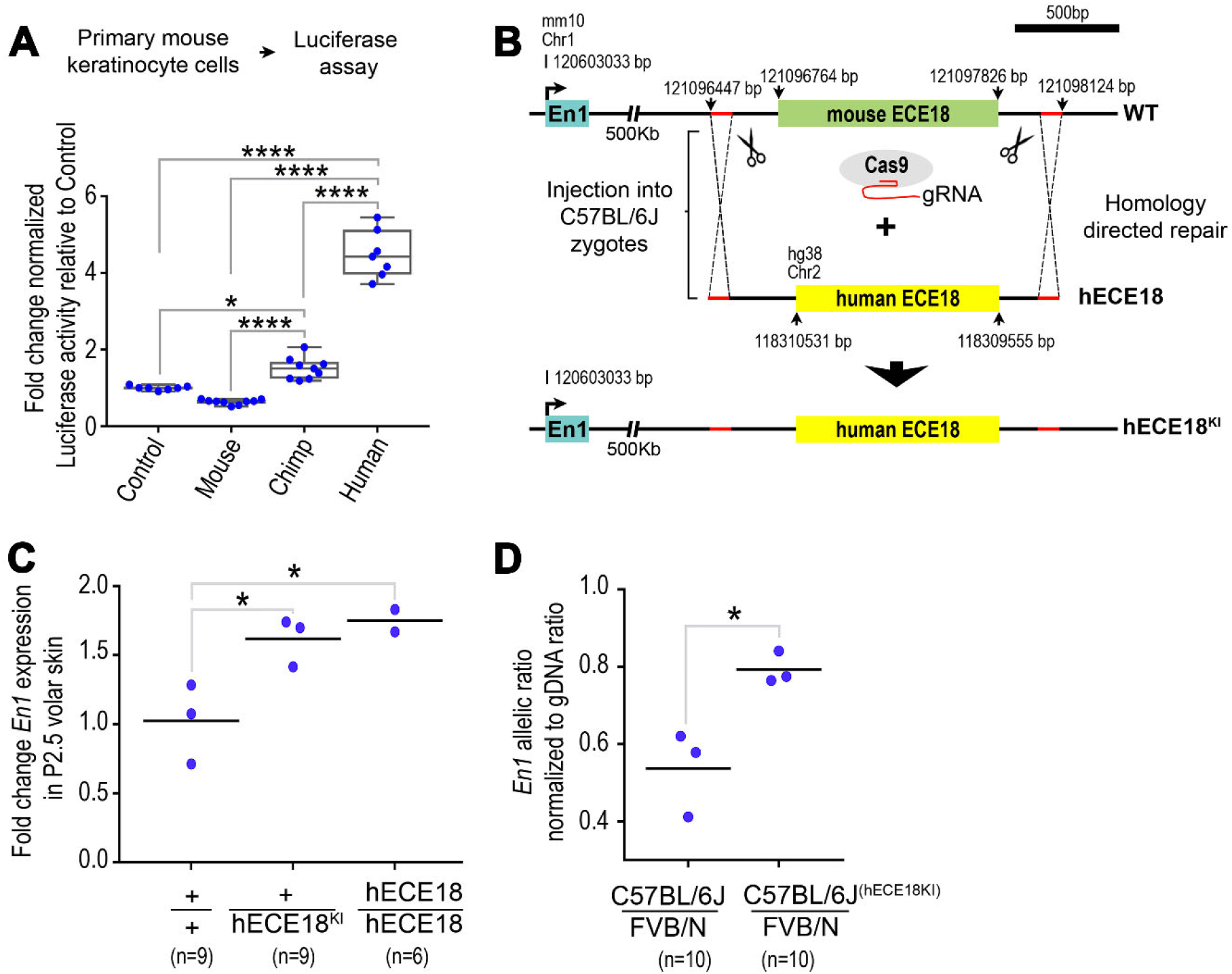
Human ECE18 acts as a developmental enhancer of *Engrailed 1* during eccrine gland development. **(A)** Fold change luciferase relative to empty reporter vector (Control) induction by ECE18 orthologs in primary mouse keratinocytes. Each point represents an individual biological replicate. Median (line), 25%-75% percentiles (box bounds) and min and max (whiskers) are shown. **(B)** Strategy to generate hECE18 knock-in (hECE18^KI^) mouse model. **(C)** Fold change in *En1* mRNA by qRT-PCR in P2.5 volar forelimb skin of wildtype (+/+), hECE18^KI^ heterozygote (hECE18^KI^ /+) and hECE18^KI^ homozygote (hECE18^KI^/hECE18^KI^) mice relative to wildtype. Each point represents a single biological sample of pooled volar skin from both forelimbs of three mice. Genotype mean is shown as a line. **(D)** Normalized ratio of C57BL/6J : FVB/N *En1* allelic expression in volar forelimb skin of wildtype (C57BL/6J / FVB/N) and hECE18^KI^ (C57BL/6J^(hECE18KI)^ / FVB/N) hybrid mice. Ratios normalized to genomic DNA allelic ratio. Genotype mean (line). Each point represents a biological sample consisting of pooled P2.5 volar skins from both forelimbs of at least three mice. (+) wildtype allele. (KI) knock-in. The total number of animals analyzed per genotype (n). Significance assessed by one-way ANOVA and Tukey-adjusted *P*-values are reported (****P<0.0001, * P<0.05). The total number of animals analyzed per genotype (n).

To investigate the functionality of hECE18 during eccrine gland development, we generated a humanized hECE18 knock-in (hECE18^KI^) by replacing the endogenous ECE18 enhancer of C57BL/6J mice with its human ortholog (Figure 5B and S5). hECE18^KI^ mice were born at expected Mendelian ratios, were viable and fertile. Quantitative RT-PCR analysis of *En1* expression in P2.5 volar skin revealed that hECE18^KI^ heterozygotes and homozygotes had on average a 1.6-fold increase in *En1* relative to wildtype sibling controls (Figure 5C). Allele-specific analysis of *En1* transcription from the volar skin of FVB/N:C57BL/6J^wt^ and FVB/N:C57BL/6J^hECE18KI^ F1 hybrids revealed that hECE18-mediated upregulation of *En1* occurs in *cis*, consistent with hECE18 acting as an enhancer of ectodermal *En1* in developing mouse volar skin (Figure 5D).

Since hECE18 exerts a positive effect on ectodermal *En1* levels, we examined if this enhancer could promote the specification of mouse eccrine glands. We did not observe a change in eccrine gland number in hECE18^KI^ mice (Figure S6A). This may reflect the relative attenuation of hECE18 potency in the mouse. Alternatively, since our data indicate that ECE18 is not required for *En1* regulation in mouse skin, the mouse may be insensitive to the human enhancer under endogenous conditions. Accordingly, we sensitized the number of volar eccrine glands formed to *En1* levels by crossing in a single copy of the *En1* knock-out allele (En1^KO^)(Kimmel et al., 2000) and quantified the number of eccrine glands in the IFP of adult mice (Figure 6A). Consistent with our previous studies (Aldea et al., 2019; Kamberov et al., 2015), En1^KO^ heterozygotes had fewer IFP eccrine glands than wildtype controls (Figure S6B). Strikingly, we found that hECE18 could partially rescue the En1^KO^ eccrine gland phenotype since En1^KO^/+; hECE18^KI^/+ mice carrying a copy of the human enhancer had on average 17% more IFP eccrine glands than En1^KO^ alone animals (En1^KO/+^; +/+) (Figure 6A, B). This result not only shows that the hECE18 enhancer can function in a developmental context to promote the formation of more eccrine glands, but also implicates the upregulation of ectodermal *En1* levels as the underlying mechanism for this effect.

**Figure 6.**
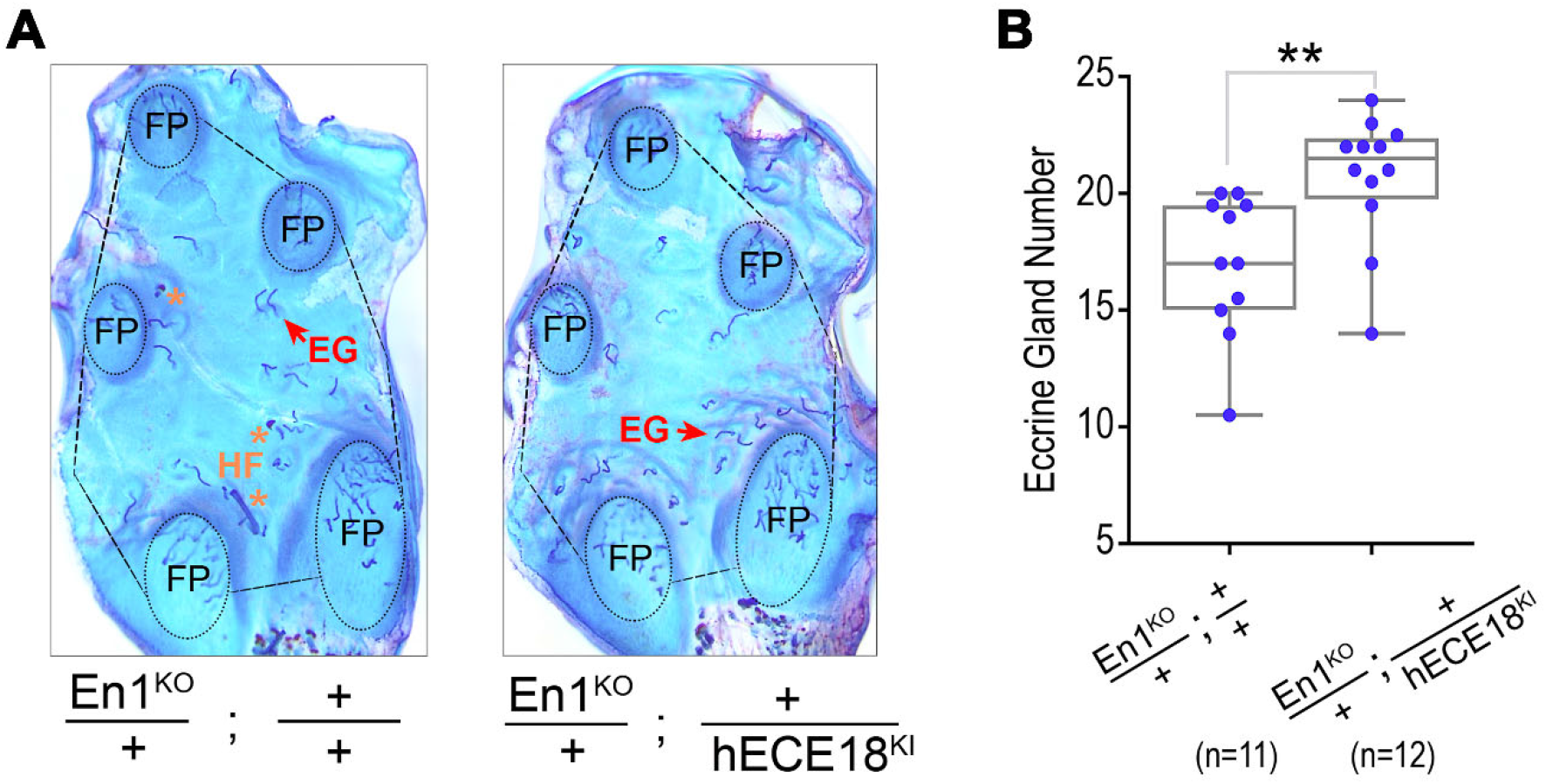
Human ECE18 upregulates *En1* to promote eccrine gland formation in mouse skin. **(A)** Representative stained epidermal preparations of volar forelimb skin from En1^KO^/+; +/+ and En1^KO^/+; +/hECE18^KI^ adult mice. The number of eccrine glands in the interfootpad area (IFP, outlined), and excluding the footpads (FP, circled) were quantified in analyses in (g). Hair follicle (HF, *), eccrine gland (EG). **(B)** Quantification of IFP eccrine glands in En1KO/+; +/+ and En1KO/+; +/hECE18^KI^ mice. Each point represents the average number of eccrine glands in the IFP across both forelimbs of a mouse. Median (line), 25%-75% percentiles (box bounds) and min and max (whiskers) are plotted. Significance assessed by a two-tailed T-test and ** *P*<0.01. (+) wildtype allele. (KO) knock-out. (KI) knock-in. The total number of animals analyzed per genotype (n).

## DISCUSSION

Natural selection has shaped many unique human attributes but the genetic bases of most of these features are unknown. This study uncovers a genetic and developmental basis for the evolution of one of the defining physiological adaptations of humans. That human ECE18 not only upregulates *EN1* expression in human skin cells, but also in a developmental model of eccrine gland formation, provides compelling evidence for a similar functionality for this enhancer in humans. Combined with the finding that human-specific mutations have increased the relative strength of hECE18, our study supports an evolutionary scenario in which a stronger hECE18 produced a relative upregulation of ectodermal *EN1* during development and thus a concomitant increase in the number of eccrine glands specified in human skin relative to that of other apes (Figure 7).

**Figure 7.**
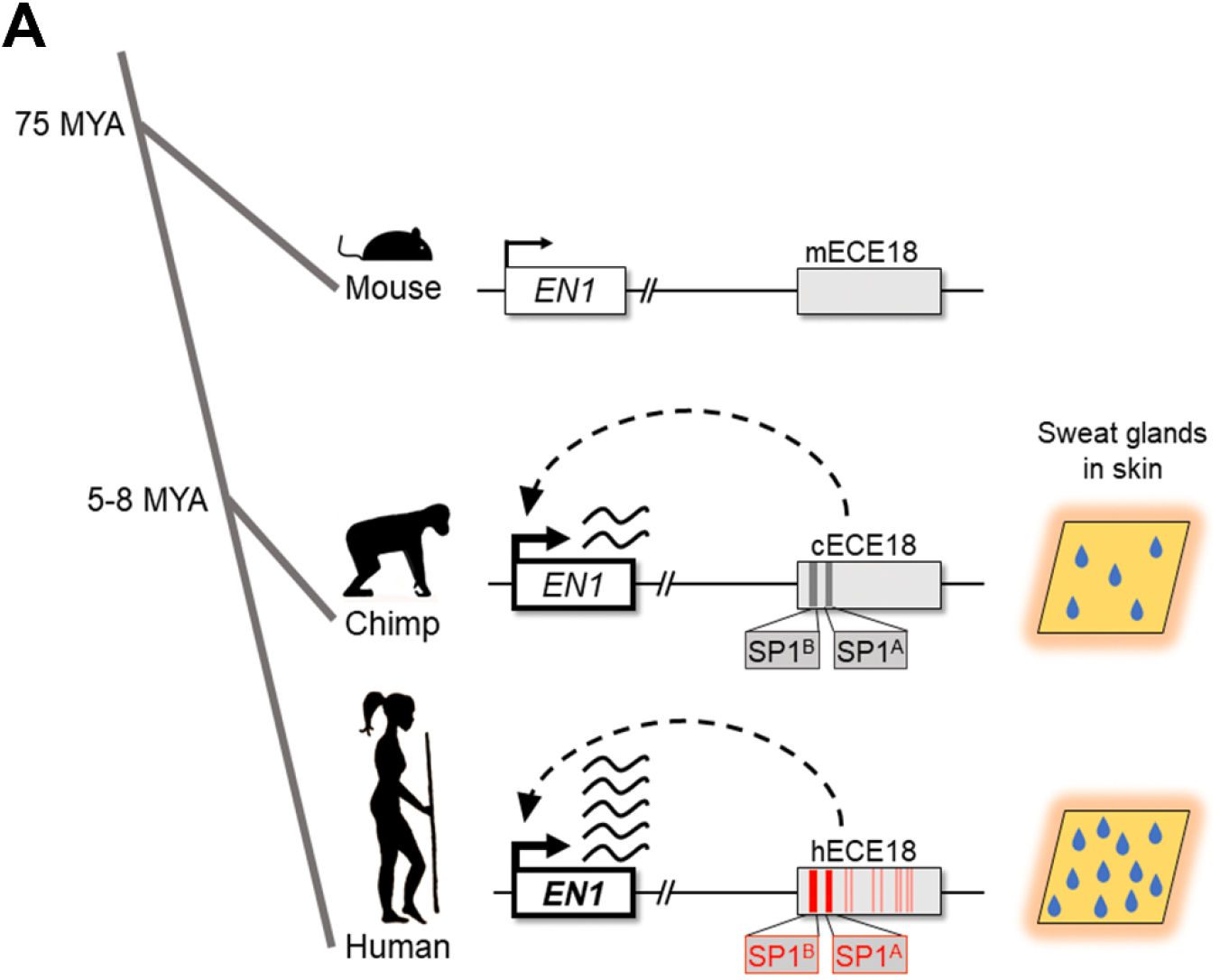
A genetic basis for the evolution of increased eccrine gland density in human skin. **(A)** ECE18 is a developmental enhancer of the anthropoid *Engrailed 1* locus. Over the course of hominin evolution, the recurrent mutation of ECE18, particularly at the SP1 sites SP1^A^ and SP1^B^ (highlighted in bold), coordinately enhanced ECE18 enhancer strength to increase the number of eccrine glands specified in human skin by potentiating developmental *EN1* expression. Derived human substitutions relative to other apes underlying higher activity of hECE18 are in red.

An intriguing feature of hECE18 is that its potency is the product of the interaction of multiple point mutations scattered throughout the enhancer. Derived human base-pair substitutions at conserved primate SP1 binding sites SP1^A^ and SP1^B^ appear to have played the greatest individual role in increasing hECE18 activity. Intriguingly, five of the remaining derived substitutions which contribute to increased hECE18 activity lie within the human accelerated region 2xHAR20. While future additional work will be required to tease apart the epistatic interactions between the mutations within and outside the HAR, and to unravel the specific steps the HAR played in the evolution of this critical trait, it is worth noting that our findings directly implicate a bioinformatically defined HAR in the evolution of an adaptive human trait.

That hECE18 gains evolved in a stepwise manner raises the possibility that the eccrine gland density of *Homo sapiens* evolved gradually, on par with the level of hECE18 activity. To understand this, we must parse out the order in which hECE18 variants evolved and the threshold at which hECE18 activity gains affect *EN1* and eccrine phenotypes. In terms of considering evolutionary mechanisms more broadly, our findings imply that in addition to the traditional paradigm whereby adaptive traits evolve as a consequence of modifications to multiple components of a developmental program (Carroll, 2003), recurrent mutation of a single functional element that incrementally alters its functionality is also a mechanism to produce gradual change in a trait over time.

A final point to consider is whether the evolution of hECE18 can solely explain humankind’s ten-fold increase in eccrine gland density over that of other apes. That hECE18 produced a subtle increase in eccrine gland number in the mouse knock-in may be reflective of the attenuated potency of this enhancer in mouse skin. Determining whether there are associations between variation in eccrine gland density and the handful of rare hECE18 polymorphisms in modern humans will shed light on the relative importance of hECE18 in controlling this human trait (Figure S7). Alternatively, there may be additional loci that contributed to the evolution of humans’ elaborated eccrine gland density. This would be consistent with the finding that eccrine gland number is controlled by multiple loci, including *En1*, in mice (Kamberov et al., 2015). Indeed, we have previously shown that variation at another locus, *EDAR*, is responsible for population-level differences in eccrine gland density among modern humans (Kamberov et al., 2013). Distinguishing between these possibilities will require comprehensive delineation of the molecular pathways that regulate eccrine gland development. The mechanistic insight into how hECE18 is regulated provides a foundation to interrogate these factors and to investigate the origins of other related traits, such as the generalization of eccrine glands to the non-volar skin. Moreover, because of the importance of *Engrailed 1* in the development of other organs and tissues, including derived human brain structures such as the cerebellum (Barton and Venditti, 2014; Wurst et al., 1994), our findings may shed light on differences in the modulation of human *EN1* in these contexts and thus the etiology of other evolutionarily significant human phenotypes.

## Acknowledgements

The authors thank Iain Mathieson, Pantelis Rompolas, Clifford J. Tabin, Daniel E. Lieberman, Benjamin Voigt, Christopher Brown, and Adam Aharoni for helpful discussions on this project and on the manuscript. The authors thank Constance Cepko and Clifford J. Tabin for training, technical help and support on ultrasound-guided injection. In addition, the authors thank Sylvain Lapan for providing constructs and technical training on lentiviral production and transduction protocols. Research reported in this publication was supported by a McCabe Fund Pilot Award from the Perelman School of Medicine, a National Science Foundation BCS-1847598 award and the National Institute of Arthritis Musculoskeletal and Skin Diseases of the National Institutes of Health under award number R01AR077690 to YGK. Any opinions, findings, and conclusions or recommendations expressed in this material are those of the authors and do not necessarily reflect the views of the National Science Foundation. The content is solely the responsibility of the authors and does not necessarily represent the official views of the National Institutes of Health.

## Contributions

YGK and DA conceived of the study, designed the experiments and wrote the manuscript. DA, YA, BK, SS and BW performed the experiments. YGK and DA analyzed and interpreted the data.

## Supplementary Information

**Figure S1.**
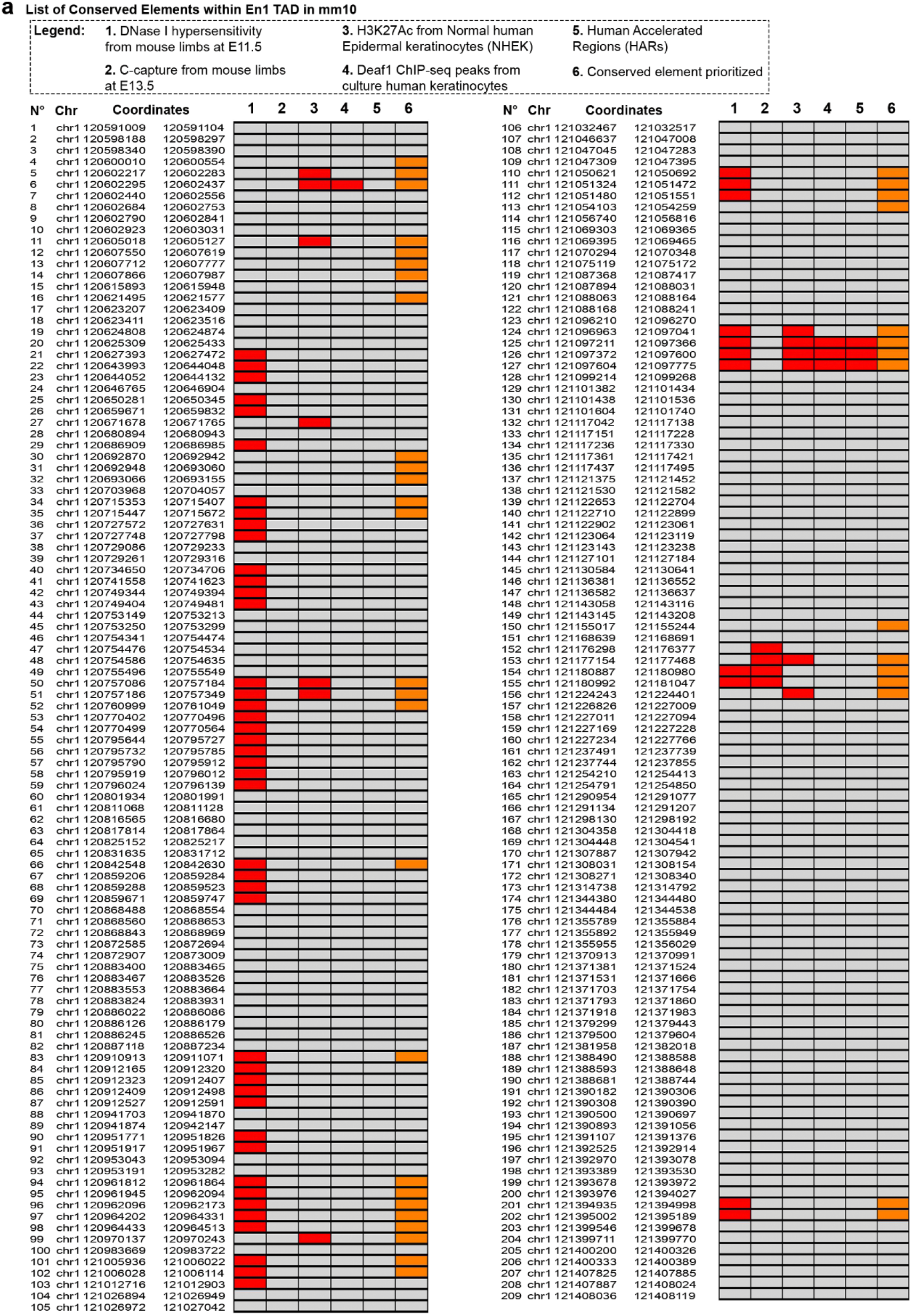
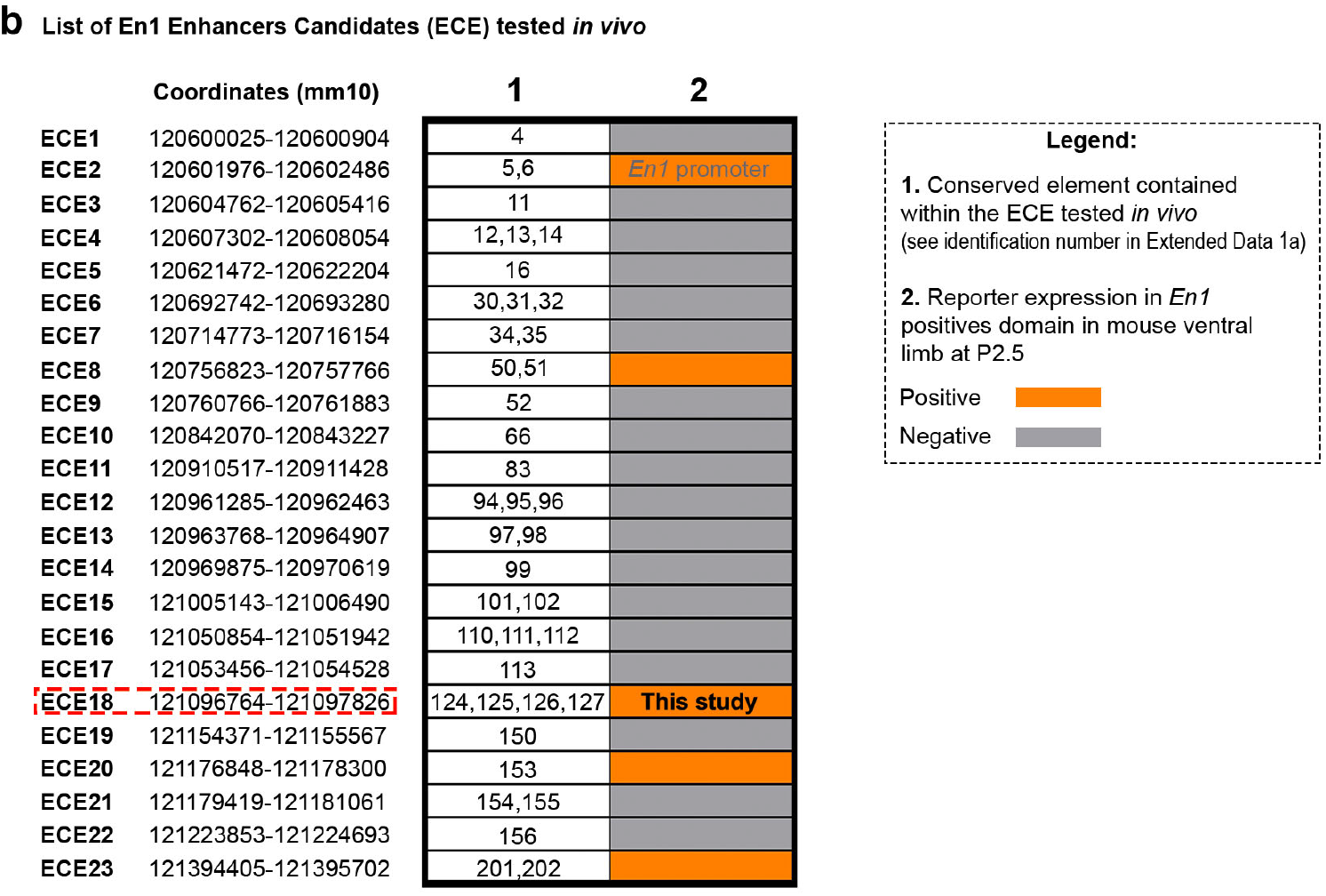
Genomic location and characteristics of conserved elements and *Engrailed 1* Candidate Enhancers (ECE). **(a)** Coordinates (mm10) of 209 conserved elements within the *EN1* TAD identified by phastCons across placental mammals. Each element has a corresponding identifier (N). Criteria used to prioritize conserved elements: overlap with published datasets of epigenomic marks associated with enhancer presence (columns 1-3) ^39,40^; overlap with DEAF1 ChIP-seq peaks^7^, which is a transcription factor we recently showed positively regulates *Engrailed 1* in human and mouse keratinocytes (column 4) ^40^; overlap with annotated human accelerated regions (HARs) (column 5) ^41-45^. Overlap is indicated in red. Prioritized conserved elements which were used as kernels for ECEs are highlighted in orange in column 6. **(b)** Genomic coordinates (mm10) of top 23 ECEs tested in mouse transgenic assays. Conserved elements (N) contained within each ECE are listed in column 1. ECEs that induced eGFP-positive clones within the *En1*-positive expression domain (basal keratinocytes of volar limb) are indicated by orange color in column 2. ECE18 is boxed in red.

**Figure S2.**
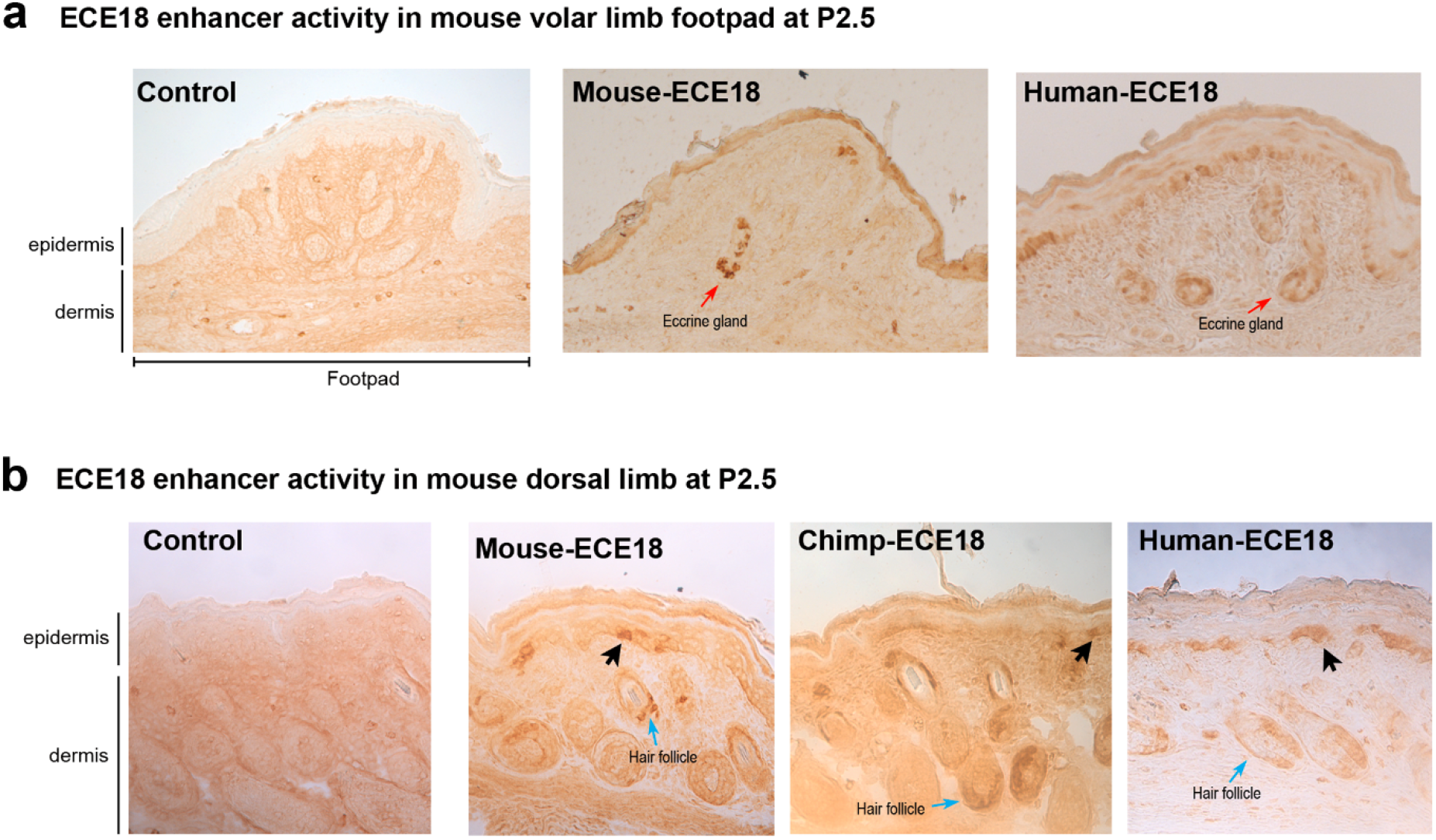
ECE18 activity in transgenic mouse skin. **(a)** Representative images of P2.5 dorsal distal autopod skin from mouse, chimpanzee and human ECE18 transgenic mice. eGFP expression was detected by anti-GFP antibody and HRP/DAB coupled immunohistochemistry. GFP positive clones (black arrow). The large hair follicles (blue arrow) which like eccrine glands derive from basal keratinocytes during development (Joost et al., 2016; Liu et al., 2013), and are characteristic of dorsal skin are also present. **(b)** Representative images of GFP antibody-stained mouse and human ECE18 transgenic skin. Cross-section of stained footpad is shown. In contrast to the IFP in which eccrine glands are being specified and still at the placode stage, the eccrine glands of the footpads (red arrow), which were specified days earlier are already undergoing differentiation as evidenced by their invagination into the dermal layer. GFP expression is visualized using HRP-DAB coupled immunohistochemistry. Control images from transgenic skin infected with lentivirus carrying minimal promoter and eGFP-reporter cassette alone.

**Figure S3.**
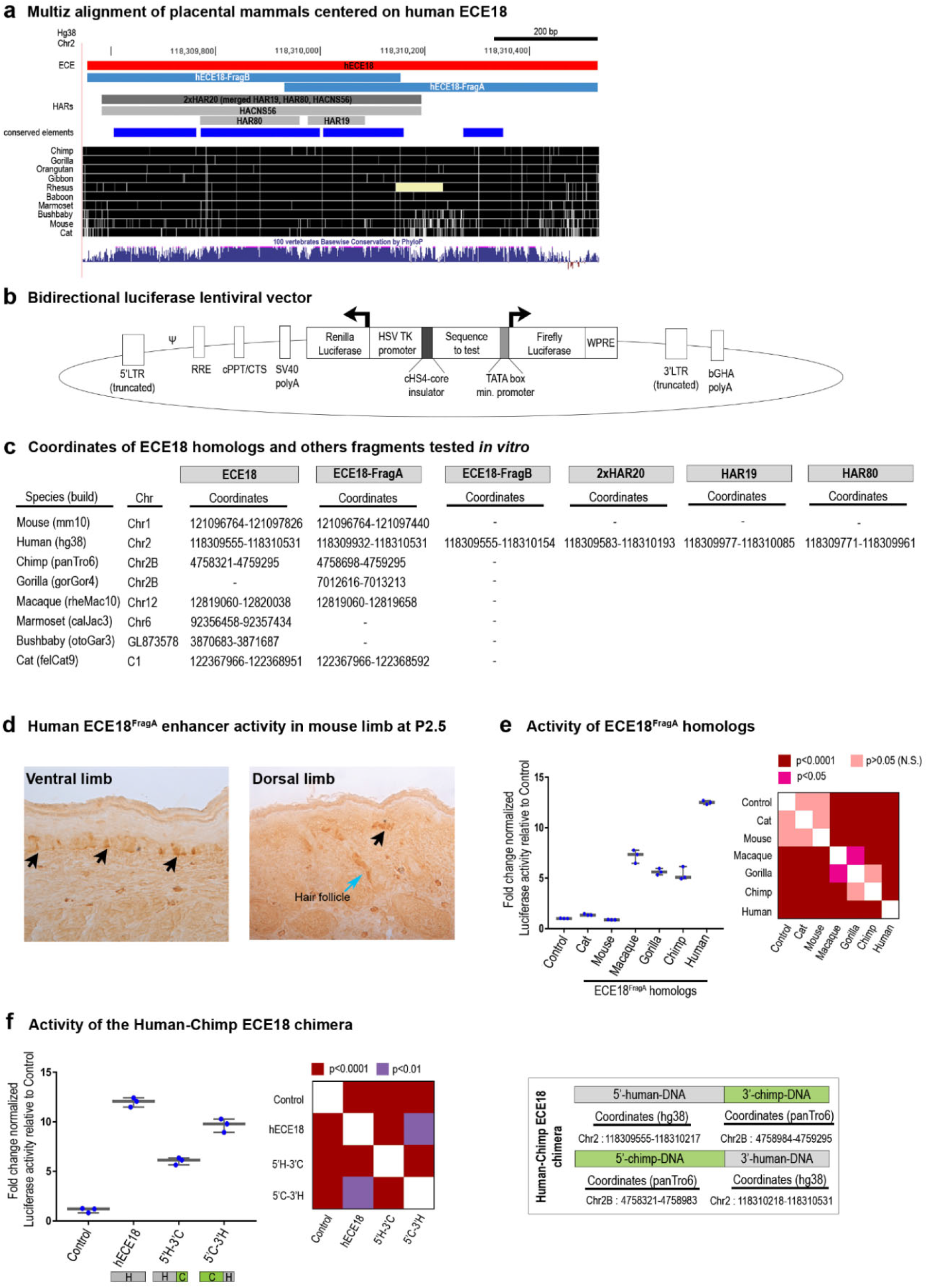
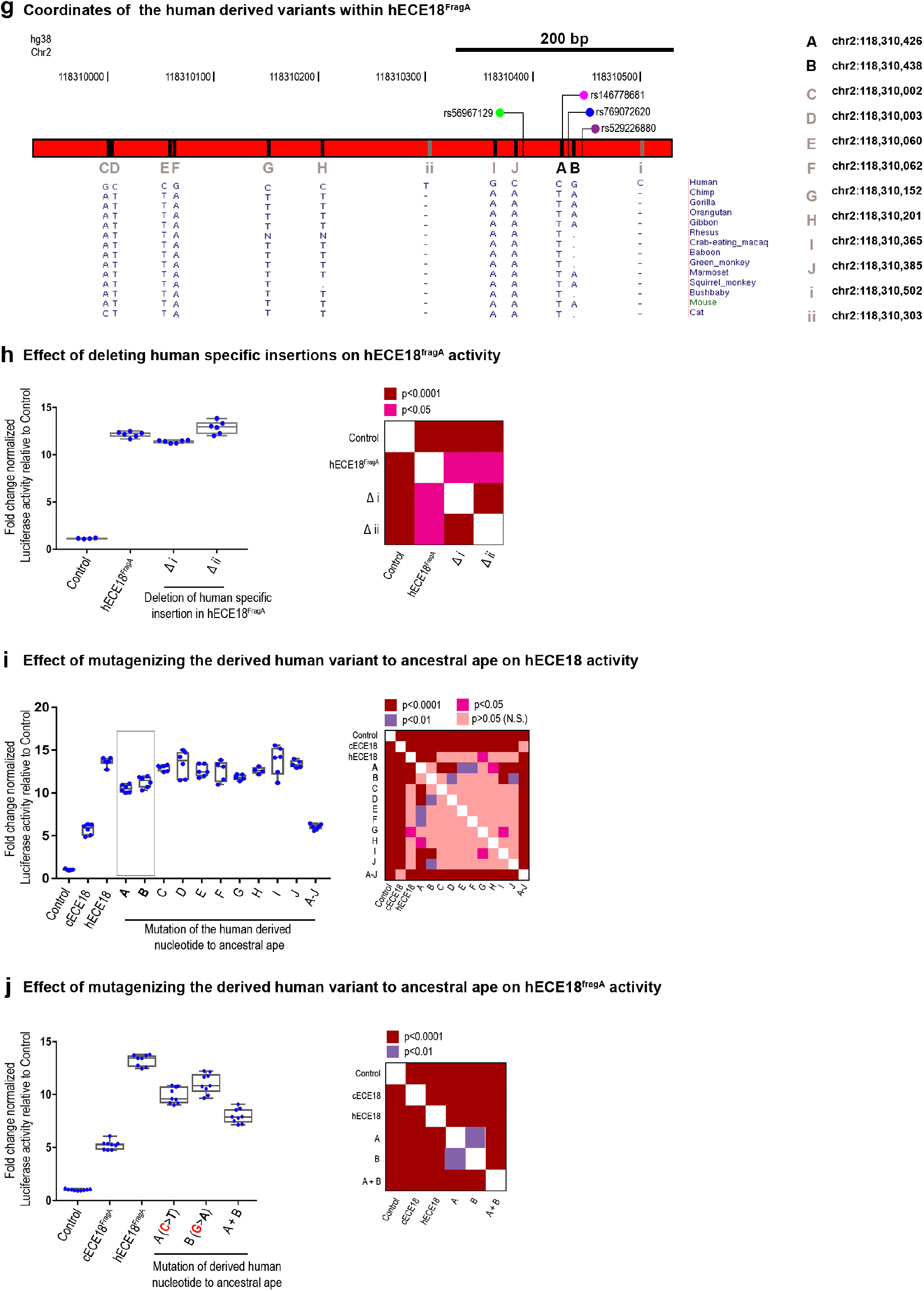
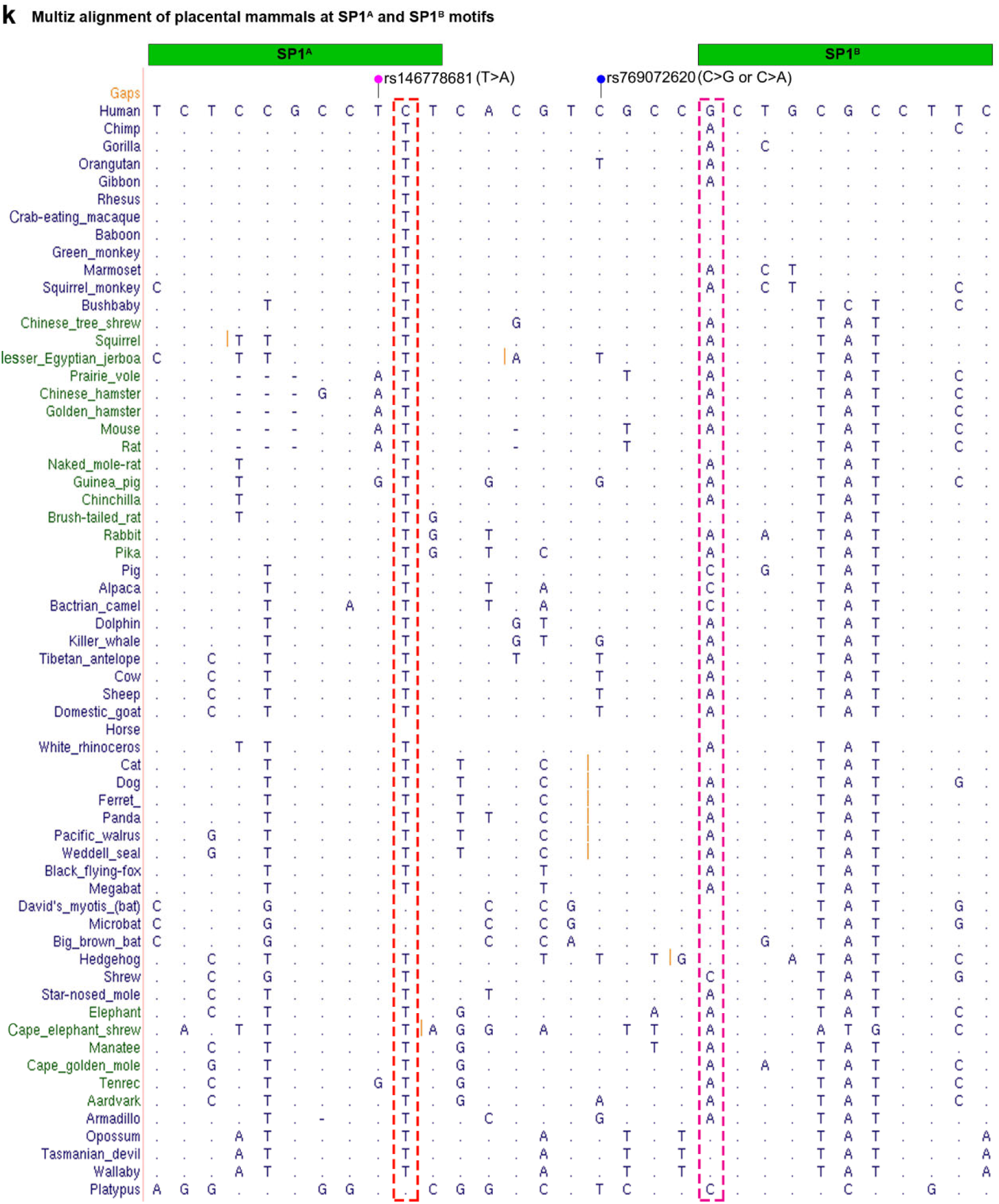
Regional dissection of ECE18 enhancer activity. **(a)** Multiz alignment of placental mammals centered on human ECE18 (hECE18). hECE18 was split into two fragments hECE18^FragA^ and hECE18^FragB^. 2xHAR20 (dark grey) is a merged element that contains HACNS56, HAR19 and HAR80 (light grey) ^42,44,49^. Conserved elements used as kernels are (#124, #125, #126 and #127) shown in dark blue. **(b)** Modified Stagia3 bidirectional luciferase lentiviral reporter vector used to test enhancer activity in cultured keratinocytes. LTR indicates long terminal repeat; Ψ, packaging signal; RRE, rev response element; cPPT, central polypurine tract; SV40 polyA, simian vacuolating virus 40 polyadenylation signal; HSV TK promoter, Herpes simplex virus thymidine kinase promoter; cHS4core, insulator core derived from the chicken CHS4 element; Sequence to test; WPRE, woodchuck posttranscriptional regulatory element; bGH polyA, bovine growth hormone polyadenylation signal. **(c)** Coordinates of ECE18 mammalian homologs and fragments tested in this study. Genome builds are indicated. **(d)** Representative images of GFP antibody stained sections from P2.5 forelimbs of hECE18^FragA^ transgenic mouse. GFP positive clones are visualized by HRP-DAB coupled immunohistochemistry so positive clones appear brown (black arrows). Dorsal hair follicle (blue arrow). **(e)** Comparative quantitative activity of mammalian ECE18^FragA^ orthologs in cultured human GMA24F1A keratinocytes. Fold change normalized luciferase activity relative to Control (empty vector) is plotted. **(f)** Fold change luciferase quantitative activity of hECE18 human: chimp chimeric enhancers relative to Control (empty vector). Maps of the human-chimp chimeric enhancers are shown (box). **(g)** Genomic location and alignments of derived human variants (A-J), derived human-specific insertions (i and ii), and modern human polymorphic variants (rs56967129, rs146778681, rs769072620, rs529226880) within hECE18^FragA^ are shown. Human coordinates in hg38. **(h)** Fold change normalized luciferase activity of hECE18^FragA^ after deletion of derived human insertions i and ii. **(i)** Effect of mutagenizing derived human bases A-J to the ancestral ape base on hECE18 enhancer activity in luciferase assays. Fold change induced by chimp ECE18 (cECE18) and hECE18^MutA-J^, in which all 10 derived bases were mutated to the ancestral ape, are also shown. **(j)** Fold change luciferase activity relative to Control (empty vector) upon mutagenesis of hECE18^FragA^ human derived variants A and B alone or in combination to ancestral ape base. **(k)** Multiz alignments of placental mammals centered on the SP1^A^ and SP1^B^ motifs. A or B human derived variants are highlighted in red and pink, respectively. Human variants rs146778681 and rs769072620 are indicated. In panels **(e, f, h, i, j)** each point represents an individual biological replicate and the median (line), 25%-75% percentiles (box bounds) and min and max (whiskers) are plotted. Significance determined by one-way *ANOVA* and Tukey’s-adjusted *P*-values are shown in heatmaps. All assays performed in cultured human GMA24F1A keratinocytes.

**Figure S4.**
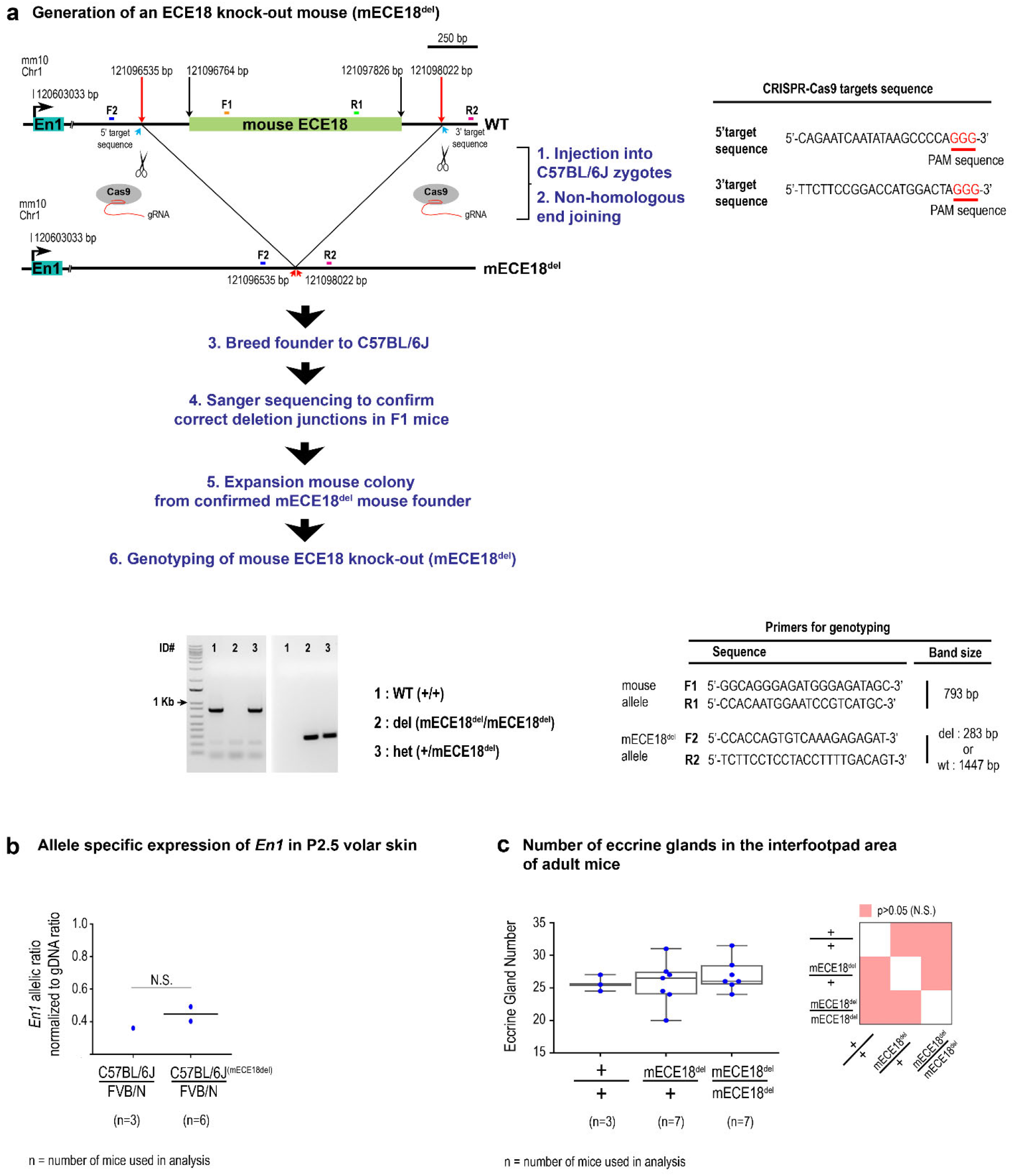
Generation and characterization of volar phenotypes of ECE18 knock-out mouse. **(a)** Generation of an ECE18 knock-out mouse (mECE18^del^) by CRISPR-Cas9 mediated genome editing. CRISPR-Cas9 target sequence and genotyping strategy are shown. Correct deletion junctions were confirmed by Sanger sequencing of F1 pups. **(b)** Normalized ratio of C57BL/6J : FVB/N allelic expression of *En1* from P2.5 volar forelimb of wildtype (C57BL/6J : FVB/N) and hECE18del (C57BL/6J^(hECE18del)^ : FVB/N) F1 hybrid mice. Ratios are normalized to the allelic ratio in F1 genomic DNA. Each point represents the mean value across three technical replicates for one or two biological samples consisting of pooled P2.5 volar skins from both forelimbs of three mice. **(c)** Quantification of the number of eccrine glands in the forelimb IFP of adult wildtype (+/+), mECE18^del^ heterozygous (+/mECE18^del^) and homozygous mECE18^del^/ mECE18^del^ mice. Each point represents the average number of IFP eccrine glands across both forelimbs of an individual mouse. The total number of animals analyzed per genotype (n). In panel (b) significance was assessed by a student’s T-test (two-tailed). In panel **(c)** significance assessed by one-way *ANOVA*. Tukey-adjusted *P*-values are shown in a heatmap and the median (line), 25%-75% percentiles (box bounds) and min and max (whiskers) are plotted. N.S., not significant.

**Figure S5.**
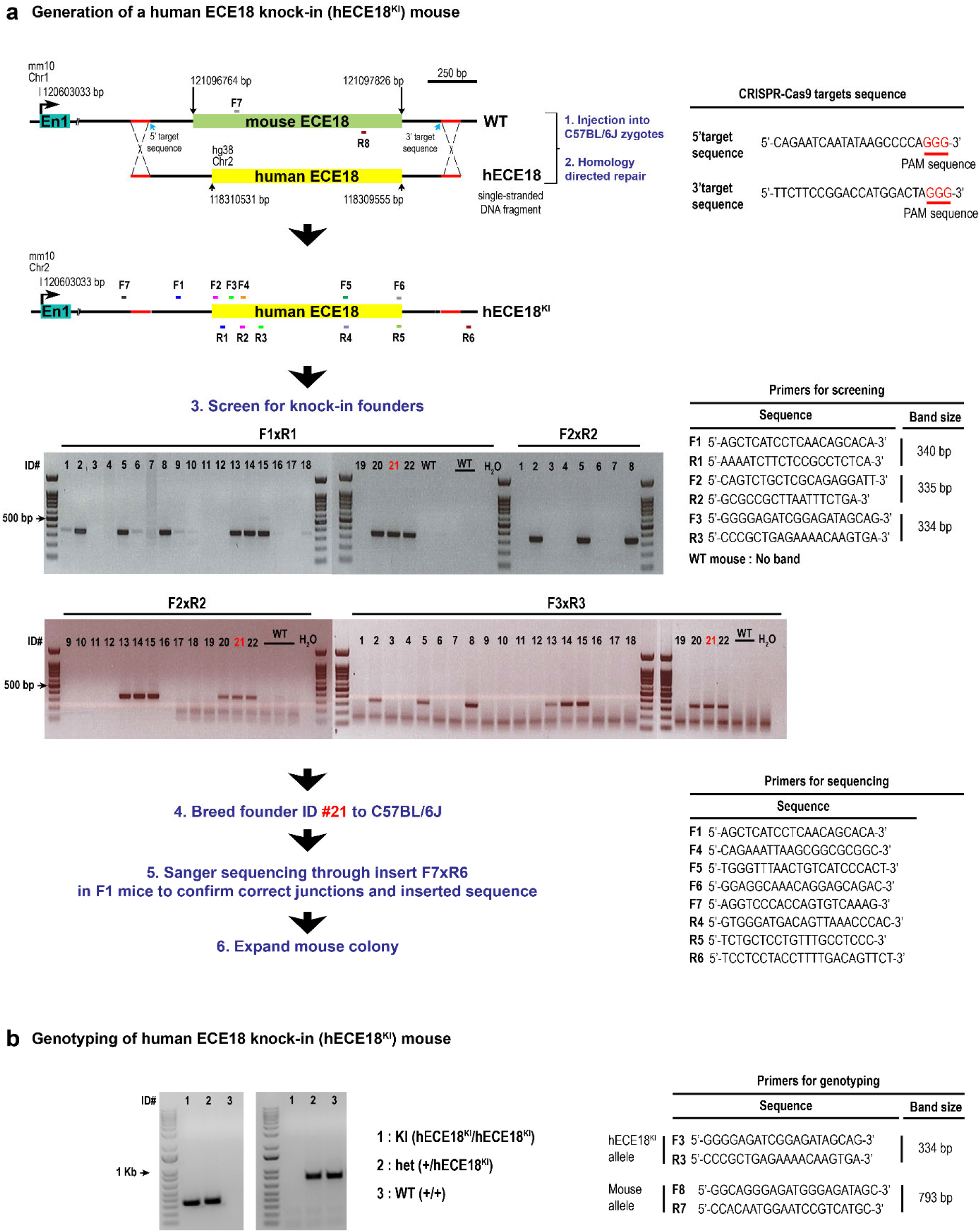
Generation of human ECE18 knock-in (hECE18^KI^) mice. **(a)** Detailed overview of the generation of a human ECE18 knock-in mouse (hECE18^KI^). CRISPR-Cas9 technology was used to replace the endogenous mouse ECE18 with the orthologous human ECE18 sequence. A single knock-in founder mouse (ID #21) was identified and bred to a C57BL/J male generate F1 pups. F1 pups were screened to confirm transmission of the knock-in allele and correct targeting and 2 pups validated to have correct junctions and insert sequence were used to generate two hECE18^KI^ lines. Each founder F1 mouse was bred onto C57BL/6J for two more generations prior to phenotypic analyses. Phenotypic analyses reported are based on progeny derived from both F1 founder lines at the N3 generation. CRISPR-Cas9 targets sequence, and primers used are listed. **(b)** Representative agarose gel for genotyping hECE18^KI^ mice. Sequences of genotyping primers used are shown.

**Figure S6.**
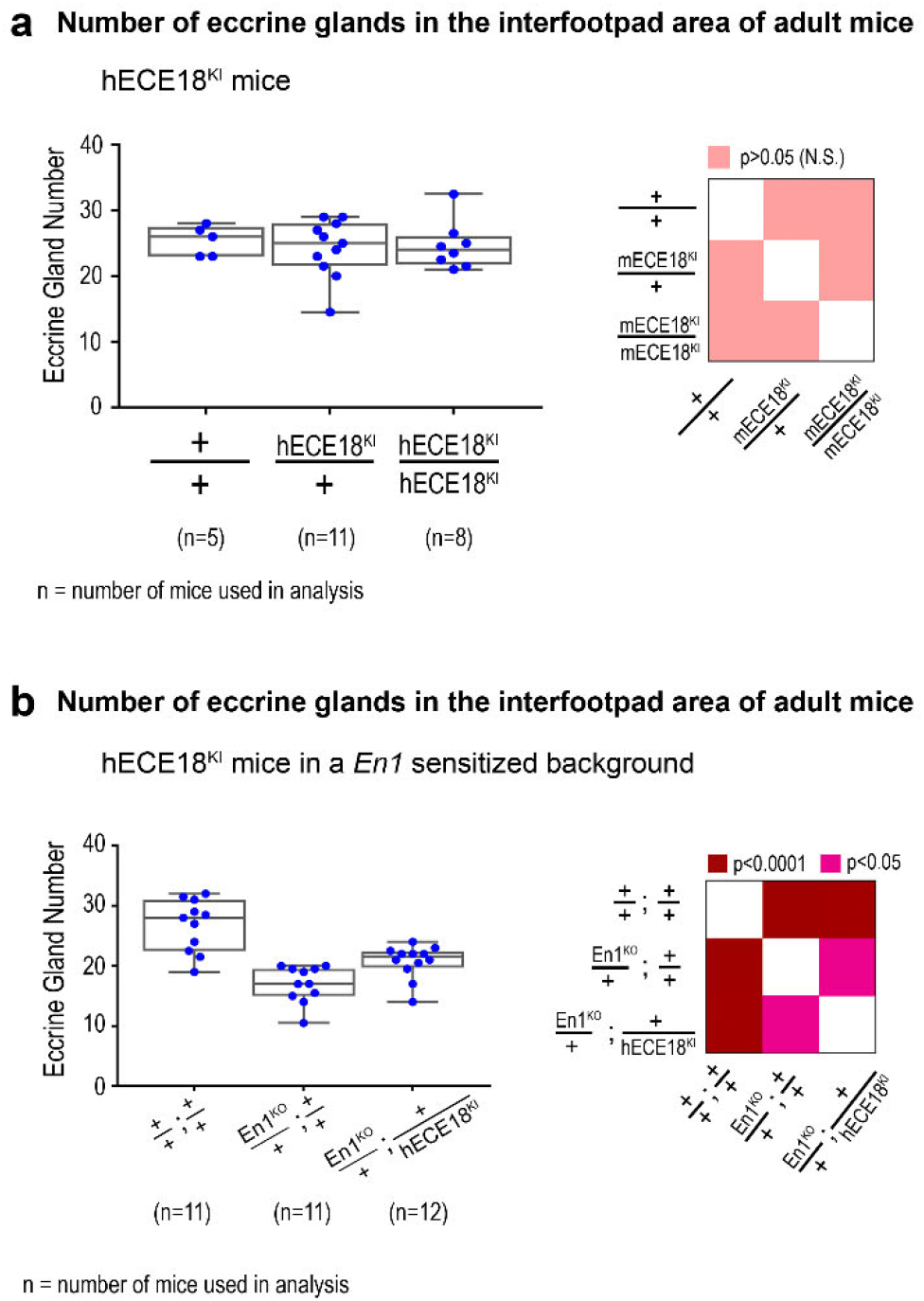
Effect of hECE18^KI^ and En1^KO^ alleles on mouse interfootpad eccrine gland number. **(a)** Effect of hECE18 on IFP eccrine gland number in a wildtype genetic background. IFP eccrine gland number in the forelimb adult wildtype (+/+), +/hECE18^KI^ and hECE18^KI^/hECE18^KI^ mice is plotted. **(b)** Effect of En1^KO^ allele on eccrine gland number. The number of eccrine glands in the forelimb IFP of adult +/+; +/+, En1^KO^/+; +/+, and En1^KO^/+; +/hECE18KI mice is plotted. Values for animals carrying En1^KO^ are also reported in the main text in Fig.3g. In panels **(a, b)** each point represents the average number of IFP eccrine glands across both forelimbs of an individual mouse and median (line), 25%-75% percentiles (box bounds) and min and max (whiskers) are plotted. The total number of animals analyzed per genotype (n). Significance was assessed by one-way *ANOVA* and Tukey-adjusted *P*-values are reported in heatmaps. +, wildtype allele. KI, knock-in. KO, knock-out. N.S., not significant.

**Figure S7.**
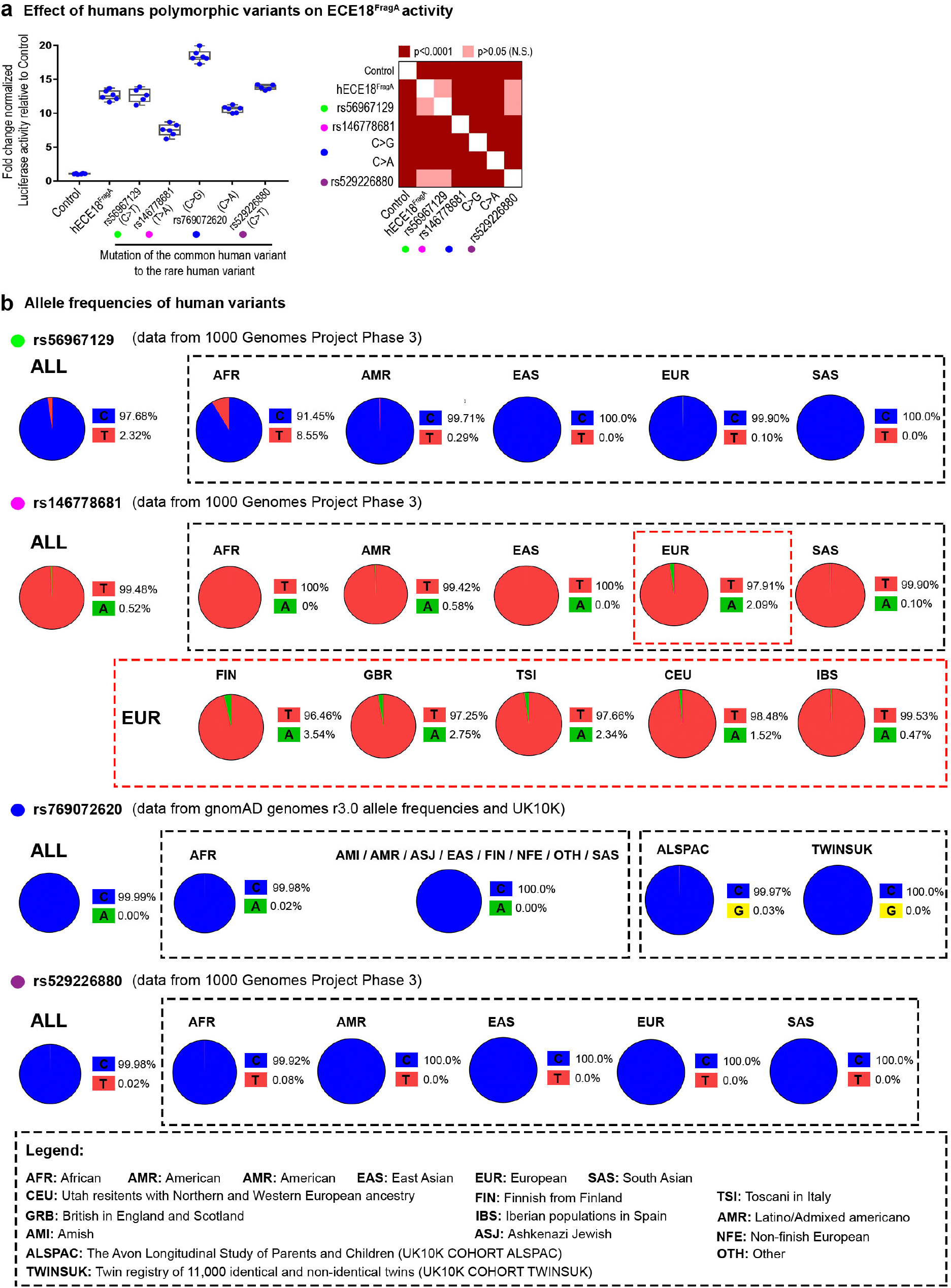
Effect of modern human single nucleotide polymorphisms on the hECE18^FragA^ enhancer activity. **(a)** Fold change normalized luciferase induction relative to Control (empty vector) by hECE18^FragA^ containing minor allele at rs56967129, rs146778681, rs769072620 and rs529226880. **(b)** Allele frequencies of polymorphic human variants rs56967129, rs146778681, rs769072620, rs529226880 are shown. Allele frequencies are from the 1000 Genomes Project Phase 3, gnomAD genomes r3.0 and UK10K datasets(1000 Genomes Project Consortium et al., 2015; UK10K Consortium et al., 2015). In panel **(a)** each point represents an individual biological replicate and the median (line), 25%-75% percentiles (box bounds) and min and max (whiskers) are plotted. Significance determined by one-way *ANOVA* and Tukey’s-adjusted *P*-values are reported in the heatmap. Assays were performed in human GMA24F1A cultured keratinocytes.

## Materials and Methods

### Identification of *Engrailed 1* candidate enhancers for *in vivo* testing

Conserved elements were called using phastCons (Pollard et al., 2010; Siepel et al., 2005) from alignment of placental mammals using the phastCons60way dataset. We restricted our scan to the *EN1* topologically associated domain (TAD) defined by Hi-C in normal human cultured keratinocytes (NHEK) (Chr2: 118000000-118880000 (hg38)) since studies on the compartmentalization of the mammalian genome suggest that elements controlling expression of *EN1*, or indeed any locus, are likely to be located within this smaller interval(Dixon et al., 2012, 2015). We called 209 conserved elements based on strings of at least 50 base pairs (bps) of consecutive high phastCons (Siepel et al., 2005) values using an empirically determined cut-off of ≥98% probability of being conserved, excluding coding exons and allowing for short gaps of missing data. We prioritized 41 of these conserved elements based on overlap with the following published epigenomic marks: DNase hypersensitivity peaks from embryonic mouse limbs (Davis et al., 2018), Capture-C peaks from mouse embryonic limbs (Andrey et al., 2017), H3K27Ac ChIP-seq peaks from mouse embryonic limbs (Davis et al., 2018), H3K27ac ChIP-seq peaks from NHEK cells (Davis et al., 2018), coordinates of known human accelerated regions (Capra et al., 2013; Pollard et al., 2006; Prabhakar et al., 2006), and Deaf1 ChIP-seq peaks (Aldea et al., 2019). Using each of these 41priority element as a kernel, we expanded these genomic regions to 1000bp-1500bp because known enhancers tend to be hundreds of base pairs long, collapsing our list to 23 *Engrailed 1* candidate enhancers (ECEs) for functional testing.

### Mice

CD1 (Crl:CD1) timed pregnant mice and FVB/NCrl mice were purchased from Charles River Laboratories. C57BL/6J mice were purchased from the Jackson Laboratories. En1^KO^ mice were generated in the lab of Dr. Alexandra Joyner (Kimmel et al., 2000) and were obtained from the laboratory of Susan Dymecki (Harvard Medical School). En1^KO^ mice were bred onto C57BL/6NTac (Taconic Biosciences) for at least 10 generations. All experiments were performed in accordance with approved IACUC protocols.

### Generation of lentiviral-mediated transient transgenic mice

The generation of lentivirus-mediated transient transgenic mice was performed as previously described (Beronja and Fuchs, 2013; Beronja et al., 2010). In brief, each ECE or ECE fragment was cloned into the Stagia3 lentiviral enhancer reporter vector, upstream of the minimal tata-box promoter and eGFP reporter cassette (Emerson and Cepko, 2011; Wang et al., 2014). Primers used for cloning are reported in **Table S1**. High-titer lentivirus for each construct was prepared in HEK293T cells according to established protocols using a second-generation packaging system (gift from Connie Cepko Harvard Medical School). Transfection of HEK293T cells was carried out using polyethylenimine (Polysciences Inc.). Following concentration, lentivirus for each tested DNA element was mixed in a 7:1 ratio with a control virus expresses mCherry under a ubiquitous promoter (modified FUtdTW in which tdTomato was replaced with an mCherry expression cassette was a gift from Connie Cepko Harvard Medical School). Expression of mCherry was used to identify successfully transduced mice at harvest. Virus for each tested element in this study, mixed with FUtdTW, was injected into the amniotic cavity of embryonic day (E) 9 CD1/NCrl mouse embryos (Charles River Labs). Injections were carried out under ultrasound guidance using the Vevo 2100 ultrasound imaging system (Visualsonics, Toronto Canada) equipped with a 35-50 MHz mechanical transducer as described previously (Beronja and Fuchs, 2013; Beronja et al., 2010). At E9, lentiviral particles are able to stably integrate into the genomes of cells of the single layer mouse ectoderm but cannot pass beyond the underlying basement membrane, allowing for ectoderm-specific generation of viral transgenic mice(Beronja et al., 2010). All layers of the skin, including the basal keratinocyte layer, derive from the single layer ectoderm at E9. At least six embryos were injected per tested DNA element. All survival surgeries were carried out in accordance with approved IACUC protocols.

On post-natal day (P) 2.5, pups were sacrificed and both forelimbs and both hindlimbs were processed for expression of eGFP in mCherry positive animals. At this stage *En1* is expressed throughout basal keratinocytes of the distal volar skin, is focally upregulated in eccrine gland placodes of the IFP and is critical for specifying IFP eccrine gland number (Figure 1A)(Kamberov et al., 2015). This spatial pattern is not recapitulated by other transcripts encoded in the TAD. Positive elements were called based on the observation of multiple eGFP positive clones in at least three individual transduced mice. Of note, since viral integration into the infected cell genome occurs independently and at random, each clone of positive cells constitutes an individual transgenic event.

### *in situ* hybridization, immunohistochemistry and imaging

Whole forelimb and hindlimb autopods (the distal segment of the limb) were embedded in OCT (Tissue Tek) and cryo-sectioned at a thickness of 10-12 μm. *En1 in situ* hybridization (ISH) was performed as previously described (Kamberov et al., 2015). HRP (Horseradish peroxidase)/DAB(3,3-diaminobenzidine) immunostaining was performed as follows. Tissue sections were fixed in 4% Paraformaldehyde (PFA) and then washed in Phosphate buffered saline (PBS). Endogenous peroxidase was blocked using 0.3% hydrogen peroxide. Tissue was blocked in PBST (PBS + 0.1% Tween) + 10% normal donkey serum before incubating in chicken anti-GFP primary antibody (1:2000, Jackson). After washing, samples were incubated with biotin-SP-conjugated rabbit anti-chicken secondary antibody (1:250, Jackson ImmunoResearch). Samples were washed and incubated in Vectastain ABC reagent (Vector laboratories). Enzymatic detection was carried out using the DAB peroxidase substrate kit (Vector Laboratories) according to manufacturer’s instructions. Cytokeratin 14 primary antibody (PRP155-P CK14, 1:10000), Alexa Fluor^594^(1:250, Jackson ImmunoResearch) and 4′6-diamidino-2-phenylindole (1:5000, Sigma Aldrich). Images were acquired on a Leica DM5500B microscope equipped with a Leica DEC500 camera.

### Bidirectional luciferase vector and luciferase assays

The bidirectional luciferase lentiviral reporter was built based on (Kleinovink et al., 2018; Na et al., 2010) and using Stagia3 as a backbone (**Figure S3B**). Briefly, the different ECE18 orthologs and the various fragments described in this study (**Figure S3C**) were cloned upstream of a minimal tata-box promoter and upstream of the Firefly Luciferase reporter gene into the bidirectional luciferase lentiviral vector and using the following primers (**Table S2**). Lentivirus production and skin cell transduction was carried out as previously described (Aldea et al., 2019; McNeal et al., 2015a). Cells were harvested at 72h post-transduction, and Firefly and Renilla (for normalization) Luciferase activities were measured using the Dual-luciferase reporter assay system (Promega). The assay was performed using the SpectraMax i3x (Molecular Devices). Experiments were done independently at least 3 times in biological triplicate each time. Mutagenesis of ECE18 orthologs was carried out using standard site-directed mutagenesis protocols and primers are reported in **Table S3**.

### *in silico* motif discovery analysis

SP1 DNA binding sites were first identified by looking at conserved motifs across the ECE18 sequence and using the ECR browser with default parameters (TRANSFAC professional V10.2 library) (Ovcharenko et al., 2004). Prediction of SP1 DNA binding motifs at the selected sites was confirmed using JASPAR 2018 and affinity scores were obtained from JASPAR 2018 database (Matrix ID: MA0079.3) (Khan et al., 2018).

### Cell culture, transfection and transduction

Primary mouse keratinocytes were isolated and cultured as previously described with minor modifications (Lichti et al., 2008; Pirrone et al., 2005). Briefly, CD1 P0 pups were sacrificed, then back skin was removed from the body and incubate over night with 0.25% trypsin-EDTA (Gibco) to detach the epidermis from the dermis. Next day, the epidermis was collected in a conical tube, and keratinocytes were dissociated by mechanical shearing by nutation at slow speed for 45 minutes at 4°C degrees. Cells were plated and maintained using K-SFM media (Gibco) containing 45 μM calcium, 4% of chelex treated fetal bovine serum (Hyclone), 10 ng/mL mouse epidermal growth factor (mEGF) (Corning), 50 μg/mL bovine pituitary extract (Gibco) and streptomycin/penicillin (Sigma).

GMA24F1A human keratinocytes were a gift from Dr. Howard Green (Harvard Medical School) and were previously described (Mainguy et al., 1999). GMA24F1A cells are a clonal line of human keratinocytes that expresses *EN1* and have basal keratinocyte characteristics which we confirmed by the expression of the signature basal keratin *Keratin 14* (*KRT14*) and by the expression of *EN1*.

HEK293T cells were obtained from the laboratory of Dr. Connie Cepko (Harvard Medical School).

Transfection of HEK293T cells was carried out using polyethylenimine (Polysciences Inc.). We used a second-generation packaging system to generate the lentiviruses used in this study (gift from Connie Cepko Harvard Medical School). All viruses were produced in HEK293T cells according to established protocols. Transduction of GMA24F1A cells and primary mouse keratinocytes was carried out as previously described (Aldea et al., 2019; McNeal et al., 2015b). Packaging plasmid psPAX2 was a gift from Didier Trono (Addgene plasmid # 12260), envelope plasmid pCL-VSV was a gift from Connie Cepko (Harvard Medical School)

### Chromatin immunoprecipitation and qPCR

ChIP-qPCR was performed as previously described (Aldea et al., 2019). In brief, GMA24F1A cells were cultured in 150 mm dishes and harvested for immunoprecipitation using the EZ-Magma ChIP HiSens kit (EMD Millipore) according to the manufacturer’s instructions. DNA was cross-linked by adding formaldehyde (1% final concentration) for 10 minutes at room temperature. After cell lysis, the chromatin was sheared using a Covaris M220 sonicator and time optimized to get chromatin fragments between 100bp and 500bp. Immunoprecipitation was carried out using 1/125 SP1 Antibody (ab13370, Abcam) or 1ug of normal rabbit IgG antibody (Sigma Aldrich). Input and immune-precipitated samples were purified using a Qiagen PCR purification kit. Quantitative PCR was done using Power SYBR PCR master mix (Thermo Fisher). SP1 occupancy of a given region after the immunoprecipitation, was determined as the percentage of enrichment over the input using the following equation 100 × 2Ct^(input) − Ct(IP)^ and represented in all graphs as enrichment of SP1 or IgG. Primers used for qPCR are reported in **Table S4**. ChIP-qPCR experiment was performed 3 times and each reported value represents the average across three technical qPCR replicates for that biological sample in a single experiment.

### Generation and maintenance of mouse ECE18 knock-out (mECE18^del^) and human ECE18 knock-in (hECE18^KI^) mice

For specific details of targeting design, strategy and validation, screening of founder animals and establishment of mutant lines see **Figure S4** (mECE18^del^) and **S5** (hECE18^KI^). In brief, pairs of guide RNAs (gRNAs) were designed using the online tool http://crispr.mit.edu/. Guides were tested to generate deletion of mouse ECE18 *in vitro* in NIH3T3 cells. For the generation of hECE18^KI^, single-stranded repair template containing hECE18 genomic sequence flanked by mouse genomic sequence homology arms was synthesized, sequence-confirmed and purchased from Genewiz. The reported gRNAs which were confirmed to target mouse ECE18 *in vitro* were used to generate genome edited mice. gRNA selection, generation and in vitro testing were performed by the Perelman School of Medicine (PSOM) CRISPR Cas9 Mouse Targeting Core. Validated gRNAs along with Ca9 RNA and hECE18 repair template were microinjected into the cytoplasm of C57BL/6J one cell embryos by the PSOM Mouse Transgenic and Chimeric mouse facility. Human ECE18 knock-in fragment was confirmed by Sanger sequencing. All procedures were performed in accordance with approved IACUC protocols.

### dCas9-KRAB repression of hECE18

gRNAs were designated using the online IDT-DNA tool (https://www.idtdna.com/) (Cr1: 5′-GATTTTCATTTCCTGTGTTA-3′ and Cr2 : 5′-TCTTTTCTGTTTACCGGGGA-3′). pLV hU6-sgRNA hUbC-dCas9-KRAB-T2a-GFP, which encodes dCas9-KRAB fusion was a gift from Charles Gersbach (Addgene plasmid # 71237; http://n2t.net/addgene:71237; RRID:Addgene_71237)(Thakore et al., 2015). gRNAs were cloned into the plasmid as previously described (Thakore et al., 2015). Lentiviral production was done using HEK293T cells line, and GMA24F1A cell transduction was performed as previously described (Aldea et al., 2019) with modifications. Briefly, low confluency GMA24F1A cells were transduced. Then, cells were harvested 5 days post-transduction when between 90-100% of the cells expressed the reporter GFP protein indicating that most cells integrated the vector producing the gRNA and the dCas9-KRAB protein. Two independent experiments were carried out in biological triplicates each time. In graph (**Figure 4A**) each dot represents the average of technical triplicates for each independent biological sample.

### RNA extraction and quantitative RT-PCR

Total RNA was isolated using TRIzol (Life Technologies). Next, we clean up the RNA using the RNeasy Mini Kit (Qiagen) according to the manufacturer’s instructions. *En1* total expression was determined as previously described (Aldea et al., 2019) and using the following primers **Table S4**. Assays were performed in biological triplicates consisting of at least six pooled ventral forelimb skins from each genotype at P2.5. cDNA was generated using SuperScript III (Thermo Fisher) following the manufacturer’s instructions. quantitative RT-PCR (qRT-PCR) was done using Power SYBR PCR master mix (Thermo Fisher) and in technical triplicates each time.

### Allelic discrimination assay

*En1* allelic expression assays were performed as previously described (Kamberov et al., 2015). Briefly, ventral forelimb skin consisting of the region containing the five volar footpads and intervening IFP were dissected from P2.5 F1 C57BL/6J: FVB/N hybrid mice and RNA was extracted. Amplification of cDNA and gDNA products was done using the following primers: *En1* Forward 5′-GAGCAGCTGCAGAGACTCAA-3′ and *En1* Reverse 5′-CTCGCTCTCGTCTTTGTCCT-3′. *En1* allelic expression was determined by the relative expression of C57BL/6J (hECE18^KI^) or (mECE18^del^) vs. FVB/N as distinguished at the genotype at rs3676156. Allelic expression data was analyzed using the sequencing based QSVanalyzer software (Carr et al., 2009). Sequencing was carried out using the aforementioned *En1* Forward primer in technical triplicate for each sample. cDNA was obtained from biological triplicates consisting of six or eight pooled ventral forelimb skins for each genotype.

### Quantification of eccrine glands in forelimb interfootpad skin

Quantification of mouse IFP eccrine gland number was performed as previously described (Kamberov et al., 2013). In brief, three to four-week old mice were euthanized and the mouse ventral forelimb skin was dissected for dissociation in Dispase II (Roche) to isolate epidermal whole mount preparations. Eccrine gland ducts remain attached to the epidermis after Dispase II treatment allowing quantification of eccrine gland number based on the number of ducts emanating from the epidermis. Whole mount epidermal preparations were stained with Nile Blue (Sigma-Aldrich) and Oil Red O (Sigma-Aldrich) to visualize volar skin appendages (hair follicles and eccrine glands) and hair follicle-associated sebaceous glands, respectively. IFP eccrine gland number was quantified by averaging the number of glands across the left and right forelimb in epidermal preparations from each animal.

### Statistical analysis

Statistical analysis was performed using either ordinary one-way *ANOVA* followed by Tukey multiple comparisons correction with a single pooled variance on the means for each dataset or student’s unpaired T-test (two-tailed) using GraphPad Prism version 7.04 for Windows, GraphPad Software, La Jolla California USA, www.graphpad.com.

---

**Table S1:**
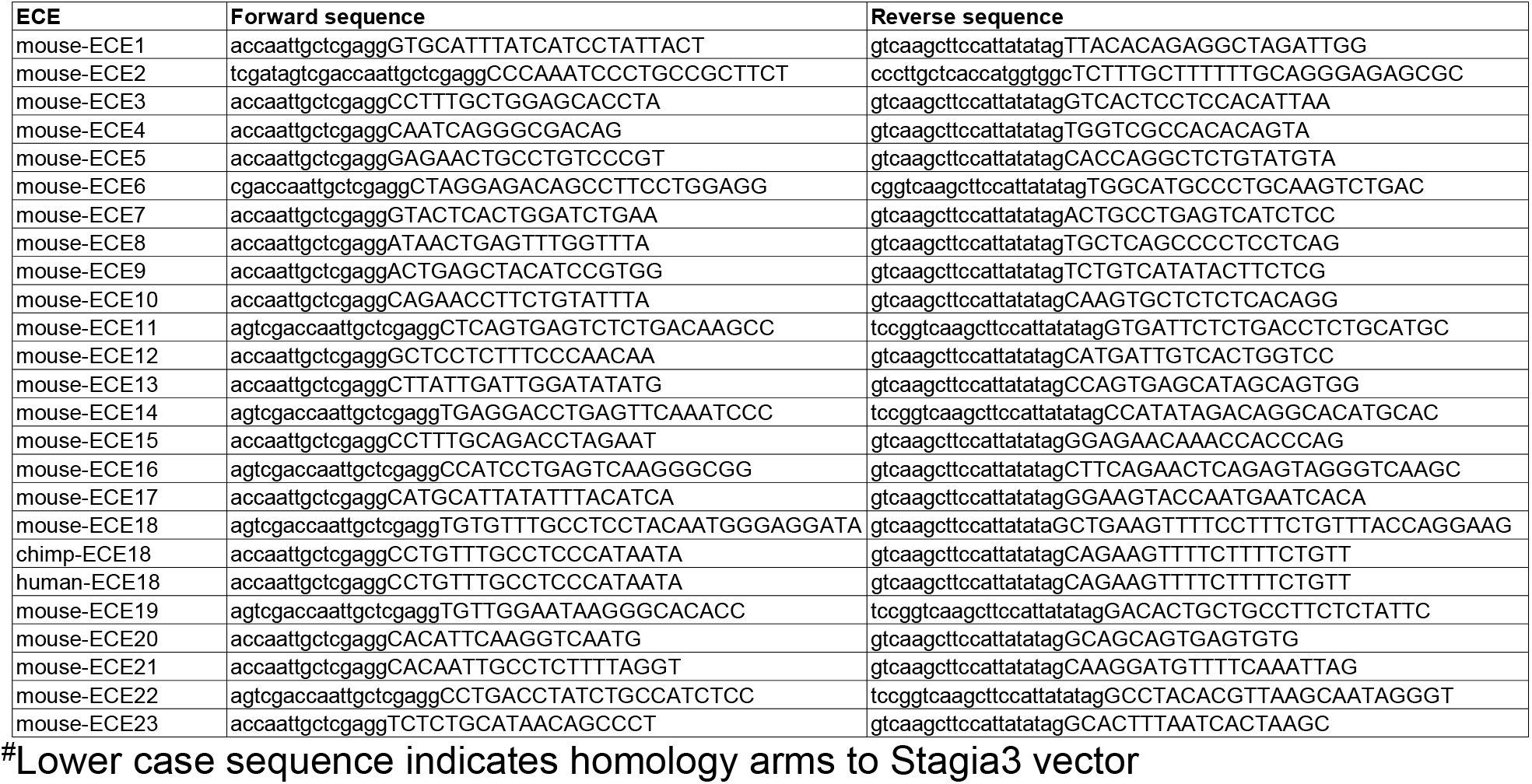
Primers used to subclone ECEs in mouse transgenic assays^#^

**Table S2:**
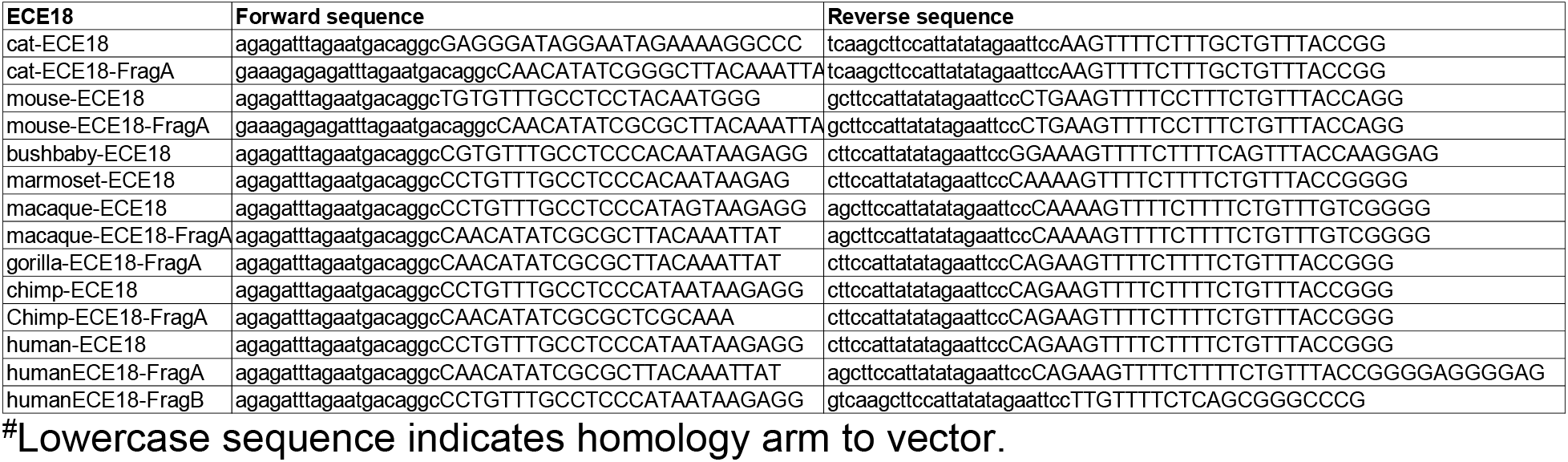
Primers used to clone ECE18 orthologs into bidirectional luciferase reporter vector^#^

**Table S3:**
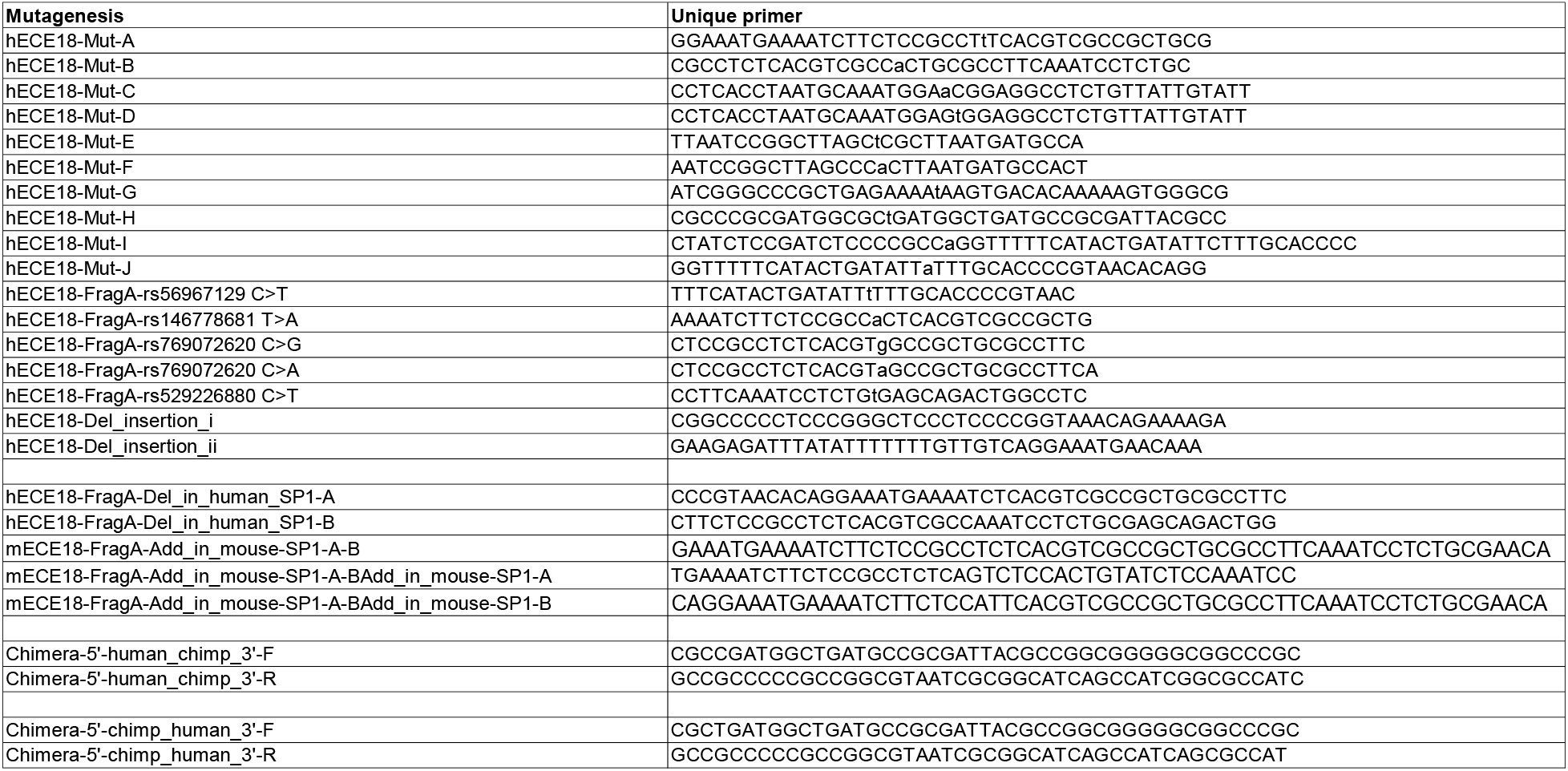
Primers used mutagenesis of ECE18

**Table S4:**
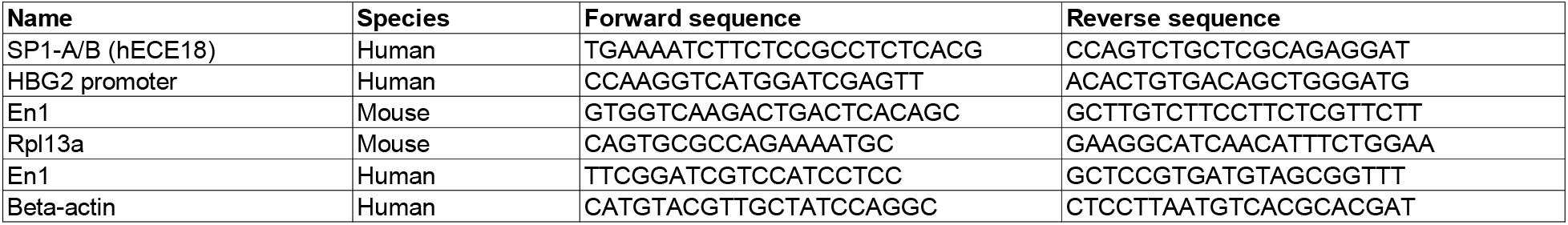
ChIP-qPCR and qRT-PCR primer sequences

## References

1. Adelman, S., Taylor, C.R., and Heglund, N.C. (1975). Sweating on paws and palms: what is its function? Am. J. Physiol. 229, 1400–1402.

2. Aldea, D., Kokalari, B., Luckhart, C., Aharoni, A., Albert, P.R., and Kamberov, Y.G. (2019). The Transcription Factor Deaf1 Modulates Engrailed-1 Expression to Regulate Skin Appendage Fate. J. Invest. Dermatol. 139, 2378–2381.e4.

3. Al-Ramamneh, D., Gerken, M., Gerken, D.M., and Riek, A. (2011). Effect of shearing on water turnover and thermobiological variables in German Blackhead mutton sheep. J. Anim. Sci. 89, 4294–4304.

4. Barton, R.A., and Venditti, C. (2014). Rapid Evolution of the Cerebellum in Humans and Other Great Apes. Curr. Biol. 24, 2440–2444.

5. Beronja, S., Livshits, G., Williams, S., and Fuchs, E. (2010). Rapid functional dissection of genetic networks via tissue-specific transduction and RNAi in mouse embryos. Nat. Med. 16, 821–827.

6. Capra, J.A., Erwin, G.D., McKinsey, G., Rubenstein, J.L.R., and Pollard, K.S. (2013). Many human accelerated regions are developmental enhancers. Philos. Trans. R. Soc. B Biol. Sci. 368.

7. Carrier, D.R. (1984). The Energetic Paradox of Human Running and Hominid Evolution. Curr. Anthropol. 25, 483–495.

8. Carroll, S.B. (2003). Genetics and the making of Homo sapiens. Nature 422, 849–857.

9. Dixon, J.R., Selvaraj, S., Yue, F., Kim, A., Li, Y., Shen, Y., Hu, M., Liu, J.S., and Ren, B. (2012). Topological Domains in Mammalian Genomes Identified by Analysis of Chromatin Interactions. Nature 485, 376–380.

10. Dixon, J.R., Jung, I., Selvaraj, S., Shen, Y., Antosiewicz-Bourget, J.E., Lee, A.Y., Ye, Z., Kim, A., Rajagopal, N., Xie, W., et al. (2015). Chromatin architecture reorganization during stem cell differentiation. Nature 518, 331–336.

11. Emerson, M.M., and Cepko, C.L. (2011). Identification of a retina-specific Otx2 enhancer element active in immature developing photoreceptors. Dev. Biol. 360, 241–255.

12. Folk, G.E., and Semken, H.A. (1991). The evolution of sweat glands. Int. J. Biometeorol. 35, 180–186.

13. Hiley, P.G. (1976). The thermoreculatory responses of the galago (Galago crassicaudatus), the baboon (Papio cynocephalus) and the chimpanzee (Pan stayrus) to heat stress. J. Physiol. 254, 657–671.

14. Kamberov, Y.G., Wang, S., Tan, J., Gerbault, P., Wark, A., Tan, L., Yang, Y., Li, S., Tang, K., Chen, H., et al. (2013). Modeling Recent Human Evolution in Mice by Expression of a Selected EDAR Variant. Cell 152, 691–702.

15. Kamberov, Y.G., Karlsson, E.K., Kamberova, G.L., Lieberman, D.E., Sabeti, P.C., Morgan, B.A., and Tabin, C.J. (2015). A genetic basis of variation in eccrine sweat gland and hair follicle density. Proc. Natl. Acad. Sci. 112, 9932–9937.

16. Kamberov, Y.G., Guhan, S.M., DeMarchis, A., Jiang, J., Wright, S.S., Morgan, B.A., Sabeti, P.C., Tabin, C.J., and Lieberman, D.E. (2018). Comparative evidence for the independent evolution of hair and sweat gland traits in primates. J. Hum. Evol. 125, 99–105.

17. Kimmel, R.A., Turnbull, D.H., Blanquet, V., Wurst, W., Loomis, C.A., and Joyner, A.L. (2000). Two lineage boundaries coordinate vertebrate apical ectodermal ridge formation. Genes Dev. 14, 1377–1389.

18. King, M.C., and Wilson, A.C. (1975). Evolution at two levels in humans and chimpanzees. Science 188, 107–116.

19. Kuno, Y. (1956). Human perspiration (Springfield, IL: Thomas).

20. Lieberman, D.E. (2015). Human locomotion and heat loss: an evolutionary perspective. Compr. Physiol. 5, 99–117.

21. Loomis, C.A., Harris, E., Michaud, J., Wurst, W., Hanks, M., and Joyner, A.L. (1996). The mouse Engrailed-1 gene and ventral limb patterning. Nature 382, 360–363.

22. Lu, C.P., Polak, L., Keyes, B.E., and Fuchs, E. (2016). Spatiotemporal antagonism in mesenchymal-epithelial signaling in sweat versus hair fate decision. Science 354, aah6102.

23. Mainguy, G., Ernø, H., Montesinos, M.L., Lesaffre, B., Wurst, W., Volovitch, M., and Prochiantz, A. (1999). Regulation of epidermal bullous pemphigoid antigen 1 (BPAG1) synthesis by homeoprotein transcription factors. J. Invest. Dermatol. 113, 643–650.

24. Montagna, W. (1963). Phylogenetic Significance of the Skin of Man. Arch. Dermatol. 88, 1–19.

25. Montagna, W. (1972). The Skin of Nonhuman Primates. Am. Zool. 12, 109–124.

26. Newman, R.W. (1970). Why Man Is Such a Sweaty and Thirsty Naked Animal: A Speculative Review. Hum. Biol. 42, 12–27.

27. Pollard, K.S., Salama, S.R., King, B., Kern, A.D., Dreszer, T., Katzman, S., Siepel, A., Pedersen, J.S., Bejerano, G., Baertsch, R., et al. (2006). Forces Shaping the Fastest Evolving Regions in the Human Genome. PLOS Genet. 2, e168.

28. Prabhakar, S., Noonan, J.P., Pääbo, S., and Rubin, E.M. (2006). Accelerated evolution of conserved noncoding sequences in humans. Science 314, 786.

29. Siepel, A., Bejerano, G., Pedersen, J.S., Hinrichs, A.S., Hou, M., Rosenbloom, K., Clawson, H., Spieth, J., Hillier, L.W., Richards, S., et al. (2005). Evolutionarily conserved elements in vertebrate, insect, worm, and yeast genomes. Genome Res. 15, 1034–1050.

30. Wang, S., Sengel, C., Emerson, M.M., and Cepko, C.L. (2014). A Gene Regulatory Network Controls the Binary Fate Decision of Rod and Bipolar Cells in the Vertebrate Retina. Dev. Cell 30, 513–527.

31. William Montagna, and Paul F. Parakhal (1974). The Structure and Function of Skin - 3rd Edition (Elsevier Inc.).

32. Wright, J.T., Grange, D.K., and Richter, M.K. (1993). Hypohidrotic Ectodermal Dysplasia. In GeneReviews(®), R.A. Pagon, M.P. Adam, H.H. Ardinger, S.E. Wallace, A. Amemiya, L.J. Bean, T.D. Bird, N. Ledbetter, H.C. Mefford, R.J. Smith, et al., eds. (Seattle (WA): University of Washington, Seattle), p.

33. Wurst, W., Auerbach, A.B., and Joyner, A.L. (1994). Multiple developmental defects in Engrailed-1 mutant mice: an early mid-hindbrain deletion and patterning defects in forelimbs and sternum. Dev. Camb. Engl. 120, 2065–2075.

## Supplementary References

1. 1000 Genomes Project Consortium, Auton, A., Brooks, L.D., Durbin, R.M., Garrison, E.P., Kang, H.M., Korbel, J.O., Marchini, J.L., McCarthy, S., McVean, G.A., et al. (2015). A global reference for human genetic variation. Nature 526, 68–74.

3. Andrey, G., Schöpflin, R., Jerković, I., Heinrich, V., Ibrahim, D.M., Paliou, C., Hochradel, M., Timmermann, B., Haas, S., Vingron, M., et al. (2017). Characterization of hundreds of regulatory landscapes in developing limbs reveals two regimes of chromatin folding. Genome Res. 27, 223–233.

4. Beronja, S., and Fuchs, E. (2013). RNAi-mediated gene function analysis in skin. Methods Mol. Biol. Clifton NJ 961, 351–361.

7. Carr, I.M., Robinson, J.I., Dimitriou, R., Markham, A.F., Morgan, A.W., and Bonthron, D.T. (2009). Inferring relative proportions of DNA variants from sequencing electropherograms. Bioinforma. Oxf. Engl. 25, 3244–3250.

8. Davis, C.A., Hitz, B.C., Sloan, C.A., Chan, E.T., Davidson, J.M., Gabdank, I., Hilton, J.A., Jain, K., Baymuradov, U.K., Narayanan, A.K., et al. (2018). The Encyclopedia of DNA elements (ENCODE): data portal update. Nucleic Acids Res. 46, D794–D801.

12. Joost, S., Zeisel, A., Jacob, T., Sun, X., La Manno, G., Lönnerberg, P., Linnarsson, S., and Kasper, M. (2016). Single-Cell Transcriptomics Reveals that Differentiation and Spatial Signatures Shape Epidermal and Hair Follicle Heterogeneity. Cell Syst. 3, 221–237.e9.

13. Kamberov, Y.G., Wang, S., Tan, J., Gerbault, P., Wark, A., Tan, L., Yang, Y., Li, S., Tang, K., Chen, H., et al. (2013). Modeling Recent Human Evolution in Mice by Expression of a Selected EDAR Variant. Cell 152, 691–702.

14. Kamberov, Y.G., Karlsson, E.K., Kamberova, G.L., Lieberman, D.E., Sabeti, P.C., Morgan, B.A., and Tabin, C.J. (2015). A genetic basis of variation in eccrine sweat gland and hair follicle density. Proc. Natl. Acad. Sci. 112, 9932–9937.

15. Khan, A., Fornes, O., Stigliani, A., Gheorghe, M., Castro-Mondragon, J.A., van der Lee, R., Bessy, A., Chèneby, J., Kulkarni, S.R., Tan, G., et al. (2018). JASPAR 2018: update of the open-access database of transcription factor binding profiles and its web framework. Nucleic Acids Res. 46, D260–D266.

16. Kimmel, R.A., Turnbull, D.H., Blanquet, V., Wurst, W., Loomis, C.A., and Joyner, A.L. (2000). Two lineage boundaries coordinate vertebrate apical ectodermal ridge formation. Genes Dev. 14, 1377–1389.

17. Kleinovink, J.W., Mezzanotte, L., Zambito, G., Fransen, M.F., Cruz, L.J., Verbeek, J.S., Chan, A., Ossendorp, F., and Löwik, C. (2018). A Dual-Color Bioluminescence Reporter Mouse for Simultaneous in vivo Imaging of T Cell Localization and Function. Front. Immunol. 9, 3097.

18. Lichti, U., Anders, J., and Yuspa, S.H. (2008). Isolation and short-term culture of primary keratinocytes, hair follicle populations and dermal cells from newborn mice and keratinocytes from adult mice for in vitro analysis and for grafting to immunodeficient mice. Nat. Protoc. 3, 799–810.

19. Liu, S., Zhang, H., and Duan, E. (2013). Epidermal development in mammals: key regulators, signals from beneath, and stem cells. Int. J. Mol. Sci. 14, 10869–10895.

20. Mainguy, G., Ernø, H., Montesinos, M.L., Lesaffre, B., Wurst, W., Volovitch, M., and Prochiantz, A. (1999). Regulation of epidermal bullous pemphigoid antigen 1 (BPAG1) synthesis by homeoprotein transcription factors. J. Invest. Dermatol. 113, 643–650.

21. McNeal, A.S., Liu, K., Nakhate, V., Natale, C.A., Duperret, E.K., Capell, B.C., Dentchev, T., Berger, S.L., Herlyn, M., Seykora, J.T., et al. (2015a). CDKN2B Loss Promotes Progression from Benign Melanocytic Nevus to Melanoma. Cancer Discov. 5, 1072–1085.

22. McNeal, A.S., Liu, K., Nakhate, V., Natale, C.A., Duperret, E.K., Capell, B.C., Dentchev, T., Berger, S.L., Herlyn, M., Seykora, J.T., et al. (2015b). CDKN2B Loss Promotes Progression from Benign Melanocytic Nevus to Melanoma. Cancer Discov. 5, 1072–1085.

23. Na, I.-K., Markley, J.C., Tsai, J.J., Yim, N.L., Beattie, B.J., Klose, A.D., Holland, A.M., Ghosh, A., Rao, U.K., Stephan, M.T., et al. (2010). Concurrent visualization of trafficking, expansion, and activation of T lymphocytes and T-cell precursors in vivo. Blood 116, e18–25.

24. Ovcharenko, I., Nobrega, M.A., Loots, G.G., and Stubbs, L. (2004). ECR Browser: a tool for visualizing and accessing data from comparisons of multiple vertebrate genomes. Nucleic Acids Res. 32, W280–286.

25. Pirrone, A., Hager, B., and Fleckman, P. (2005). Primary mouse keratinocyte culture. Methods Mol. Biol. Clifton NJ 289, 3–14.

26. Pollard, K.S., Salama, S.R., King, B., Kern, A.D., Dreszer, T., Katzman, S., Siepel, A., Pedersen, J.S., Bejerano, G., Baertsch, R., et al. (2006). Forces Shaping the Fastest Evolving Regions in the Human Genome. PLOS Genet. 2, e168.

27. Pollard, K.S., Hubisz, M.J., Rosenbloom, K.R., and Siepel, A. (2010). Detection of nonneutral substitution rates on mammalian phylogenies. Genome Res. 20, 110–121.

30. Thakore, P.I., D’Ippolito, A.M., Song, L., Safi, A., Shivakumar, N.K., Kabadi, A.M., Reddy, T.E., Crawford, G.E., and Gersbach, C.A. (2015). Highly specific epigenome editing by CRISPR-Cas9 repressors for silencing of distal regulatory elements. Nat. Methods 12, 1143–1149.

31. UK10K Consortium, Walter, K., Min, J.L., Huang, J., Crooks, L., Memari, Y., McCarthy, S., Perry, J.R.B., Xu, C., Futema, M., et al. (2015). The UK10K project identifies rare variants in health and disease. Nature 526, 82–90.

32. Wang, S., Sengel, C., Emerson, M.M., and Cepko, C.L. (2014). A gene regulatory network controls the binary fate decision of rod and bipolar cells in the vertebrate retina. Dev. Cell 30, 513–527.

